# Heteromerization of endogenous mu and delta opioid receptors tunes mu opioid receptor signaling and trafficking

**DOI:** 10.1101/455147

**Authors:** Lyes Derouiche, Muzeyyen Ugur, Florian Pierre, Anika Mann, Stéphane Doridot, Stéphane Ory, Stefan Schulz, Dominique Massotte

**Affiliations:** Centre National de la Recherche Scientifique, Université de Strasbourg, Institut des Neurosciences Cellulaires et Intégratives, Strasbourg, France; Institute of Pharmacology and Toxicology, Jena University Hospital, Friedrich Schiller University Jena, Jena, Germany

**Keywords:** G protein-coupled receptors, opioid receptor, heteromers, internalization, ERK phosphorylation, neuropathic pain, fluorescent knock-in mice, endogenous opioid peptides, hippocampus, rostral ventromedial medulla

## Abstract

Increasing evidence indicates that native mu and delta opioid receptors can associate to form heteromers in discrete brain neuronal circuits. However, little is known about their signaling and trafficking. Using double fluorescent knock-in mice, we investigated the impact of neuronal co-expression on the internalization profile of mu and delta opioid receptors in primary hippocampal cultures and *in vivo*. We established ligand selective mu-delta co-internalization upon activation by exogenous ligands and provide evidence for mu-delta co-internalization by the endogenous opioid peptide met-enkephalin, but not β-endorphin. Co-internalization was driven by the delta opioid receptor, required an active conformation of both receptors and led to sorting to the lysosomal compartment. This alteration in the mu opioid receptor intracellular fate was accompanied by sustained ERK1/2 phosphorylation. In addition, increased mu-delta neuronal co-localization in the rostral ventromedial medulla in a chronic neuropathic state suggests that mu-delta heteromers are involved in the regulation of nociceptive transmission

## Introduction

The opioid system modulates numerous physiological functions such as nociception, emotional responses, reward and motivation, cognition but also neuroendocrine physiology and autonomic functions (Gaveriaux-Ruff and Kieffer 2002, Feng et al. 2012). The system is composed of three homologous G protein-coupled receptors (GPCR) mu, delta and kappa and three endogenous peptide families the enkephalins, endorphins and dynorphins. Several decades of opioid pharmacology have uncovered the physiological complexity of the system and highlighted functional interactions across receptors. Among these, molecular cross-talk between mu and delta opioid receptors have been extensively studied in co-transfected cells in which these two receptors can physically associate to form a novel entity with specific binding and signaling properties also called heteromer (Fujita et al. 2014). Evidence for mu-delta receptor heteromers *in vivo* has also accumulated over the past years (Massotte 2015). Mapping mu and delta opioid receptors in the brain, spinal cord and dorsal root ganglia (DRG) using double fluorescent knock-in mice revealed discrete neuronal co-expression in subcortical networks essential for survival including the perception and processing of aversive stimuli with high mu-delta expression in the nociceptive pathways (Erbs et al. 2015). Moreover, electrophysiological data collected in the rostroventral medulla (RVM) established the presence of functional mu and delta opioid receptors in the same neurons (Pedersen et al. 2011). Co-immunoprecipitation in the hippocampus (Erbs et al. 2015) or the spinal cord (Gomes et al. 2004) and high affinity for MDAN-21, a bivalent ligand bridging the mu agonist oxymorphone and the delta antagonist naltrindole, suggested that the two receptors are in close physical proximity (Daniels et al. 2005). Furthermore, direct structural contact could be disrupted in the brain, spinal cord and DRGs using interfering peptides (Xie et al. 2009, He et al. 2011, Kabli et al. 2013).

Extensive literature established specific ligand binding properties, modification in signaling pathways and ligand selective opioid receptor co-internalization in heterologous systems (Fujita et al. 2014). However, our current knowledge of the ligand binding, signaling and trafficking properties of endogenous mu-delta heteromers in physiological and pathophysiological contexts remains fragmentary though crucial to understanding their functional relevance and to evaluating their physiological role(s) and potential as a therapeutic target. Devi’s research team showed cross-allosteric modulation with a positive cooperativity promoted upon binding of the first ligand in SKNSH neuroblastoma cells that naturally co-express the two opioid receptors (Gomes et al. 2000, Gomes et al. 2004, Gomes et al. 2011). In VTA slices, co-application of the delta antagonist TIPPψ together with the mu agonist DAMGO or co-application of the mu antagonist CTAP together with the delta agonists deltorphin II or DPDPE strongly suggested that mu-delta receptor heteromerization decreases opioid signal transduction (Margolis et al. 2017). In addition, constitutive recruitment of β- arrestin 2 by mu-delta heteromers was observed in SKNSH neuroblastoma cells that affected ERK1/2 signaling (Rozenfeld et al. 2007). Finally, the mu agonist DAMGO induced co-internalization and co-recycling of mu and delta opioid receptors in DRG cultures pretreated with morphine, suggesting that mu-delta heteromerization may affect the trafficking of the delta opioid receptor in these conditions (Ong et al. 2015). However, further work is now needed to fully apprehend functional changes induced by mu-delta heteromerization, in particular upon activation by endogenous opioid peptides.

In this context, we sought to better characterize the impact of mu-delta heteromerization on the intracellular fate of native mu and delta opioid receptors. We therefore took advantage of our double fluorescent knock-in mice co-expressing functional mu and delta opioid receptors respectively fused to the red fluorescent protein mCherry or the green fluorescent protein eGFP (Erbs et al. 2015) to monitor mu and delta receptor internalization in primary hippocampal cultures or *in vivo*. We show ligand specific mu-delta receptor co-internalization induced by the mu-delta biased agonist CYM51010 (Gomes et al. 2013), but also the mu agonist DAMGO and the delta agonist deltorphin II. We also demonstrate that the mu-delta heteromer sorts to the lysosomal compartment. Using the double fluorescent mouse model, the activation of mu-delta heteromers by the endogenous opioid peptide Met-enkephalin modified the intracellular fate of the mu opioid receptor and produced sustained ERK1/2 phosphorylation. Finally, we show that, in a pathological state of chronic pain, there is an increase in mu-delta receptor co-expression in the RVM, an important brain region involved in nociceptive transmission and regulation. Together, our data point to a specific physiological role of mu-delta heteromers and provide initial validation as novel therapeutic targets for innovative strategies in chronic pain management.

## Material and methods

### Animals

Double knock-in mice co-expressing fluorescent mu and delta opioid receptors (mu-mCherry / delta-eGFP) were obtained by crossing previously generated single fluorescent knock-in mice expressing delta-eGFP or mu-mCherry, as described previously (Erbs et al. 2015). Single fluorescent knock-in mice deficient for the other receptor were generated by crossing delta-eGFP with mu-knock-out mice or mu-mCherry with delta knock-out mice. The genetic background of all animals was 50:50 C57BL6/J:129svPas. Male and female adult mice (8 to 12 weeks old) were used for *in vivo* experiments.

Mice were housed in an animal facility under controlled temperature (21 ± 2 °C) and humidity (45 ± 5 %) under a 12:12 dark–light cycle with food and water ad libitum. All experiments were performed in agreement with the European legislation (directive 2010/63/EU acting on protection of laboratory animals) and received agreement from the French ministry (APAFIS 20 1503041113547 *(APAFIS#300).02).*

### Drugs

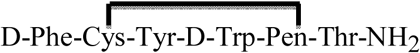 (CTAP) (C-6352), beta-funaltrexamine (β-FNA) (O-003), fentanyl citrate (F3886), naltrindole (N-2893), [D-Ala^2^, NMe-Phe^4^, Gly-ol^5^]enkephalin (DAMGO) (E-7384), Met-enkephalin (H-2016), β-endorphin (E-6361), deltorphin II (T-0658), and pertussis toxin (P-2980) were purchased from Sigma. (+)-4-[(α*R*)-α-((2*S*,5*R*)-4- Allyl-2,5-dimethyl-1-piperazinyl)-3-methoxybenzyl]-*N*,*N*-diethylbenzamide (SNC80) (cat n° 0764) was from Tocris bioscience, 1-[[4-(acetylamino)phenyl]methyl]-4-(2-phenylethyl)-4-piperidinecarboxylic acid, ethyl ester (CYM51010) (ML-335) was from Cayman chemical and Tic-deltorphin was synthesized as reported in (Salvadori et al. 1999).

#### In vivo drug administration and sample preparation

CYM51010 (10mg/kg), fentanyl (0.3mg/kg) and SNC80 (10mg/kg) were administered by i.p. injection. Met-enkephalin administration was performed by unilateral intracerebroventricular (icv) injection into the lateral ventricle of adult mice (3-4 months old) under general anasthesia. In brief, mice were anesthetized using ketamine/xylazine (100/10 mg/kg, i.p.) and placed in a stereotaxic apparatus (David Kopf, Tujunga, CA). 5µL Metenkephalin 75mM was injected at coordinates: anteroposterior −0.2 mm, ±1 mm lateral-2mm dorsoventral. For opioid receptor phosphorylation, mice were sacrificed 30 minutes after drug administration, brains collected and immediately frozen in liquid nitrogen then kept at −80° C until further use. For immunohistochemistry, mice were perfused at the indicated time points with cold paraformaldehyde 4% in phosphate buffer (PB) 0.1M pH 7.4 as described previously (Erbs et al. 2015). After 24 hours post-fixation, brain sections (30 µm thick) were cut with a vibratome (V1000 Leica) and kept floating in PB.

### Receptor phosphorylation

Frozen brain samples were transferred into ice-cold detergent buffer (50 mM Tris-HCL, pH 7.4; 150 mM NaCl; 5 mM EDTA; 10 mM NaF; 10 mM disodium pyrophosphate; 1% Nonidet P-40; 0.5% sodium deoxycholate; 0.1% SDS; containing protease and phosphatase inhibitors) and homogenized using MINILYS workplace homogenizer (Peqlab; Erlangen, Germany). Tissue homogenates were centrifuged at 16000 × g for 30 min at 4 °C after 1 h lysis at 4 °C. For the immunoprecipitation of mouse delta-eGFP or muP-mCherry, anti-GFP beads or anti-RFP beads (NanoTag; Goettingen, Germany) were used. The tissue supernatant were incubated with the beads for 2 h at 4 °C and proteins were eluted from the beads with SDS-sample buffer for 30 min at 50 °C. Proteins were separated on 7.5% SDS-polyacrylamide gels and after electroblotting, membranes were incubated with anti-pS363 (Assay Biotechnology Company Inc; CA, USA), anti-pT370, anti-S375, anti-T376 or anti-T379 antibodies at a concentration of 0.1 µg/ml and detected by chemiluminescence with ECL according to the manufacturer instructions (Doll et al. 2011, Just et al. 2013). To confirm equal loading of the gels, blots were stripped and re-probed with the phosphorylation-independent anti-GFP antibody (Synaptic Systems; Goettingen, Germany) or anti-RFP (ChromoTek GmbH; Planegg-Martinsried, Germany).

### Primary neuronal culture

Primary neuronal cultures were performed as previously described (Derouiche et al. 2018). Briefly, P0-P3 mice pups were decapitated, hippocampi were dissected and digested with papain (20 U/mL, Worthington cat. no. LS003126). Cells were plated (8-10×10^4^ cells/well) on poly-lysine (PLL, Sigma) coated coverslips in 24-well plates for immunofluorescence studies or in 12-well plates coated with poly-lysine for ERK phosphorylation studies. Cultures were maintained for 15 days *in vitro* (DIV) with half of the medium (Neurobasal A medium supplemented with 2% B27 (GIBCO, cat. no. 17504044), 2mM glutamax (GIBCO, cat. no. 35050061), 0.5mM glutamine and penicillin/streptomycin) changed every 5-7 days. Fully matured primary neurons (DIV 10 to 14) were used for all studies.

### Ex vivo drug administration and sample preparation

Met-enkephalin, β-endorphin, DAMGO, naltrindole, CTAP, deltorphin II and Ticdeltorphin were dissolved in sterile milliQ water, CYM51010 was dissolved in saline solution with DMSO (0.2% final volume) and Tween 80 (1% final volume), SNC 80 was dissolved in DMSO at 10mg/ml. Drugs were added to the culture medium of mature neurons (as 1% of the total culture volume) (12 to 15 days *in vitro*) and incubated at 37° C as indicated. Antagonists were added to the culture medium 15 minutes before agonist treatment.

For immunofluorescence studies, cultures were washed in cold 0.1 M saline phosphate buffer pH 7.4 (PBS) and fixed with 4% Paraformaldehyde in PBS. Cells were washed 3 times with cold PBS and kept at 4°C until processing.

For Western blot analyses, the medium cells were directly harvested at the end of the drug treatment in cold 100 µL/well Laemmli buffer. Cell lysates were incubated for 5 minutes at 95°C and kept at −20°C until further use.

### ERK1/2 phosphorylation

For each condition, 15 µL of sample were separated on 12% SDS-PAGE and transferred to PVDF membranes (Immobilon, Millipore). Following blocking in 5% non-fat dry milk in 50mM Tris-HCl pH8, 150mM NaCl, 0.5 % Tween 20 (TBST), membranes were incubated overnight at 4°C with monoclonal mouse mAb anti-phospho-p44/42MAPK (anti-pERK; 1:3000; Cell Signaling #9106) in blocking buffer, washed 3 times in TBST and incubated for 2 hours at room temperature with a goat anti-mouse HRP-conjugated secondary antibody (Molecular Probes, cat. n° 3412). Chemiluminescence was detected using ECL™ Prime Western Blotting Detection Reagent (Amersham Biosciences, cat. n° RPN2232) according to the manufacturer instructions. Following detection of the phosphorylated ERK forms, membranes were stripped in 100mM Glycine buffer pH 2.6 for 1h at room temperature and processed as described above for total MAPK detection with a polyclonal rabbit antibody anti-p44/42MAPK (anti-ERK1/2; 1:5000 Cell signaling, cat n° 9102) followed by goat anti-rabbit HRP-conjugated secondary antibody (1:10.000, Millipore, cat n° AP307P). For each time point, the density of ERK 1/2 phosphorylation was normalized to the density of total ERK using the ImageJ software.

### Fluorescent detection with antibodies

Primary neuronal cultures or brain section were incubated in the blocking solution PBST (PBS with 0.2 % Tween 20 (Sigma)) and 5 % normal goat serum (Sigma)) for one hour at room temperature (20-22°C) and then in primary antibodies (table 1) diluted in the blocking solution overnight at 4°C. Cells were washed three times in PBST and incubated for two hours with the secondary antibodies diluted in blocking solution (table 1). After three washes in PBST, nuclei were stained with DAPI (1µg/mL in PBS) for 5 minutes. Samples were mounted with ProLong™ Gold Antifade mounting medium (Molecular Probes) and kept at −20° protected from light until confocal imaging.

### Mu-delta neuronal co-expression in the rostroventral medulla

Neuropathic pain was induced in double fluorescent knock-in mice by cuffing the main branch of the right sciatic nerve as previously described (Benbouzid et al. 2008, Yalcin et al. 2014). Surgeries were performed under ketamine (Vibrac, Carros, France) / xylazine (Rompun, Kiel, Germany) anesthesia (100/10mg/kg, i.p.) on male and female mice. The common branch of the right sciatic nerve was exposed, and a cuff of PE-20 polyethylene tubing (Harvard Apparatus, Les Ulis, France) of standardized length (2mm) was unilaterally inserted around it (Cuff group). Sham-operated animals underwent the same surgical procedure without cuff implantation (Sham group). Mice were perfused 8 weeks later with cold paraformaldehyde 4% in PB 0.1M pH 7.4 as described previously (Erbs et al. 2015). After 24 hours post-fixation, brain were cryoprotected and sections (30 µm thick) cut with a cryostat (Microm Cryo-star HM560) and kept floating in PB for immunohistochemistry.

### Image acquisition and analysis

Confocal images were acquired (Leica SP5) using a 63x (NA 1.4) oil immersion objective and analyzed with ICY software <u>(http://icy.bioimageanalysis.org/)</u> as previously described (Derouiche et al. 2018). Briefly, quantification was performed on a single plane image from a z-stack within two sequential steps. First, the plasma membrane and cytoplasmic compartment were defined for each neuron. The spots were then detected in each channel and the amount of co-localization determined in each region of interest. The protocol used in these analyses is available online (*NewColocalizer with binary and excel output v1_batch.xml*). The extent of internalization is expressed as the ratio of membrane/cytoplasm immunoreactivity densities for each receptor or co-localized mu-delta. Co-localization of the two receptors is expressed as the percentage of co-localized mu-mCherry and delta-eGFP signals reported to the total immunoreactivity.

Quantification of neurons co-expressing mu and delta opioid receptors in the rostroventral medulla was performed using the NDP view plus software (Hamamatsu Photonics) on images from at least 5 non consecutive sections (bregma −5.5 to −6.5) on areas including the posterior raphe nuclei (pallidus, magnus, obscurus, interpositus), the parapyramidal nucleus, the alpha part of the gigantocellulat reticular nucleus and lateral paragigantocellular nucleus according to the Paxinos atlas v4.

### Statistical analysis

Statistical analyses were performed with Graphpad Prism V7 software (GraphPad, San Diego, CA). Normality of the distributions and homogeneity of the variances were checked before statistical comparison to determine appropriate tests. One-way non-parametric (Kruskal Wallis followed by Dunn’s multiple comparison test) or parametric one-way ANOVA test (followed by Dunnett’s multiple comparison test) were used to compare different experimental groups. A two-way ANOVA followed by post-hoc Tukey’s test for multiple comparisons was used for multiple factor comparisons. Results in graphs and histograms are illustrated as means ± SEM.

## Results

### Endogenous mu-delta heteromers are present at the neuronal surface under basal conditions

Previous work showed receptor expression at the surface of hippocampal neurons in basal conditions of the fluorescent knock-in mice expressing delta-eGFP and/or mu-mCherry (Scherrer 2006; Pradhan 2010;Faget 2012; Erbs 2014; Erbs 2015). As expected, under basal conditions, both mu and delta opioid receptors were predominantly located at the plasma membrane in primary hippocampal neurons as revealed by confocal imaging of mu-mCherry and delta-eGFP associated fluorescence (**Figure 1A left panel**). Quantification of the receptor density using the ICY bioimaging software (Derouiche et al. 2018) indicated that the fluorescence density at the cell surface was 2.5-fold higher compared to the cytoplasm for either receptor (**Figure 1B**). Merged images highlighted an overlay of the green and red fluorescence at the surface of the neuron and quantification of the density of receptor co-localization indicated about 20% of the total mu-mCherry or delta-eGFP signals co-localized at the plasma membrane (**Figure 1C**) and only 10% of the receptors co-localized in the cytoplasm (**Figure 1D; see also Figure S1**). Data therefore establish close physical proximity of endogenous mu and delta opioid receptors at the plasma membrane under basal conditions and suggest constitutive mu-delta heteromerization at the surface of neurons.

**Figure 1:**
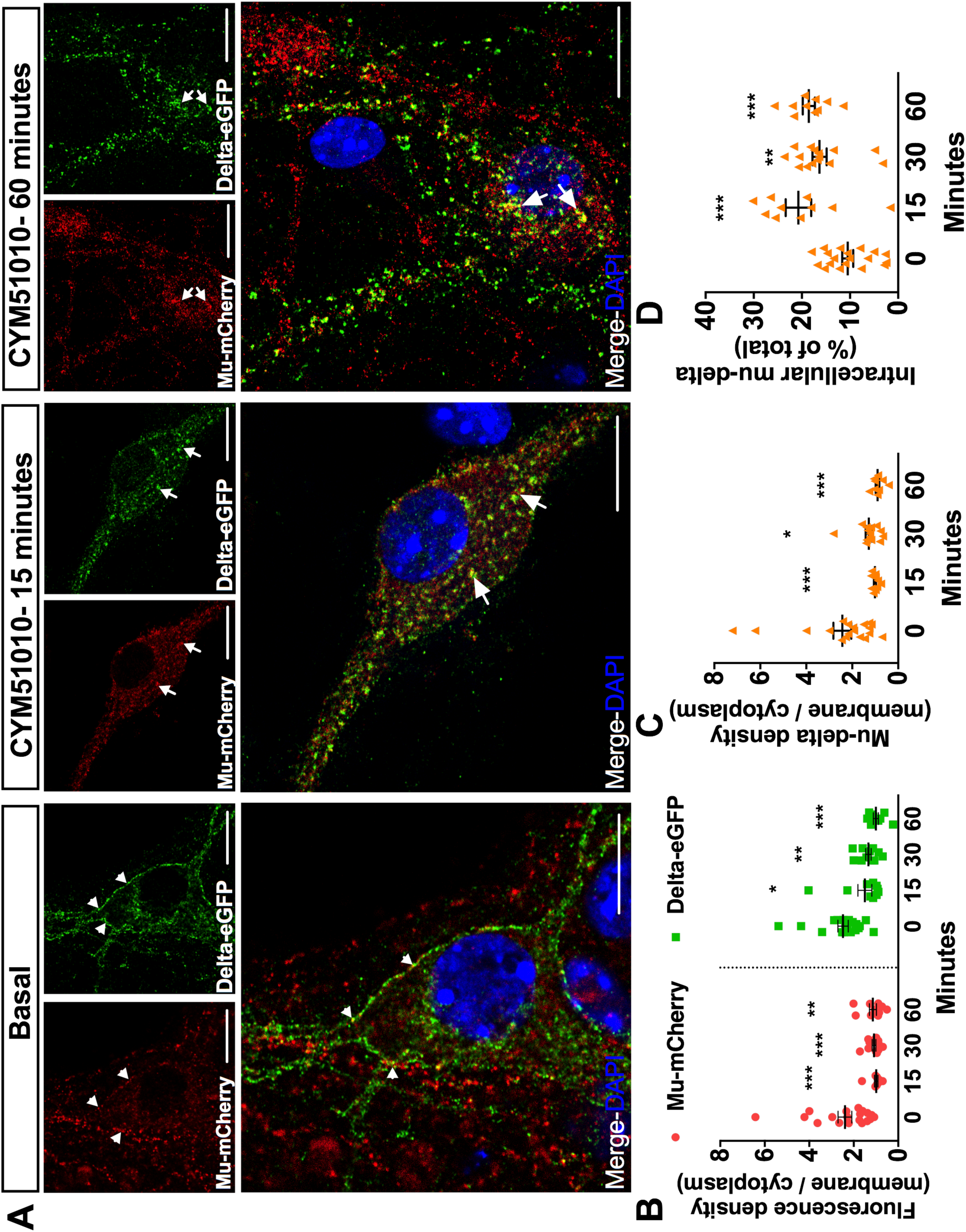
Mu and delta opioid receptors co-internalize upon CYM51010 activation in primary hippocampal cultures. **A)** Representative confocal images showing mu-mCherry and delta-eGFP fluorescence localized at the plasma membrane (arrowheads) under basal condition or internalized in vesicle-like structures 15 or 60 minutes after CYM51010 (400nM) application (arrows). Scale bar = 10µm. **B)** Receptor internalization induced by CYM51010 application (400nM) expressed as a ratio of membrane-associated versus intracellular fluorescence densities for each receptor. Two-way ANOVA F_treatment_ (3;94)=17.98 ; p-value <0.0001. F_receptor_ (1; 94)=1.06; F_interaction_ (3; 94)=0.54. Tukey’s post-hoc test for mu-mCherry ***: p <0.001; **: p = 0.01. Tukey’s post-hoc test for delta-eGFP *:p = 0.02 ; **:p = 0.002 ; ***:p < 0.001. N=10 to 20 neurons per group from at least 3 independent cultures **C)** Subcellular redistribution of mu-delta heteromers expressed as a ratio of membrane-associated versus intracellular fluorescence densities for co-localized mu-mCherry and deltaeGFP receptors. One-way ANOVA (F (3,48) = 13.64; p < 0.0001) followed with multiple comparisons Dunn’s post hoc test. *:p = 0.03 ; ***:p <0.001. N=10 to 20 neurons per group from at least 3 independent cultures. **D)** Fraction of cytoplasmic mu-delta heteromers expressed as the percentage of mumCherry and delta-eGFP overlapping objects detected in vesicle-like structures at the different time points. Kruskal Wallis test (p < 0.0001) followed with multiple comparisons Dunn’s post hoc test. **:p = 0.004 ; ***:p <0.001. N=10 to 20 neurons per group from at least 3 independent cultures.

### CYM51010 induces mu-delta receptor co-internalization and co-localization in the late endosomal compartment

CYM51010 was reported as a mu-delta biased agonist because its antinociceptive effect was blocked by an antibody selective for mu-delta heteromers and its activity was reduced in mice deficient for the mu or delta opioid receptor (Gomes et al. 2013). We therefore tested whether activation by this ligand (concentration range 10nM to 10µM) triggered mu and delta receptor internalization in primary hippocampal cultures from double fluorescent knock-in mice. CYM51010 concentrations equivalent to, or higher than, 400nM induced mu-mCherry and delta-eGFP internalization as seen from the decrease in fluorescence density associated with the plasma membrane and the appearance of fluorescent intracellular vesicles (**Figure 1A middle and right panels, Figure 1B**). Quantification of the extent of co-localization 15, 30 and 60 minutes after agonist administration showed that the fraction of mu and delta opioid receptors that co-localized at the plasma membrane significantly decreased (**Figure 1C)** whereas mu-delta receptor co-localization increased in the cytoplasm at the three time points (**Figure 1D**) establishing co-internalization of the receptors. Triple immunofluorescence labeling with LAMP1 as a marker of the late endosomal-lysosomal compartment showed increased co-localization with mu-mCherry and delta-eGFP 60 minutes after activation by CYM51010 (**Figure 2)** suggesting that mu and delta opioid receptors are targeted together to the degradation pathway.

**Figure 2:**
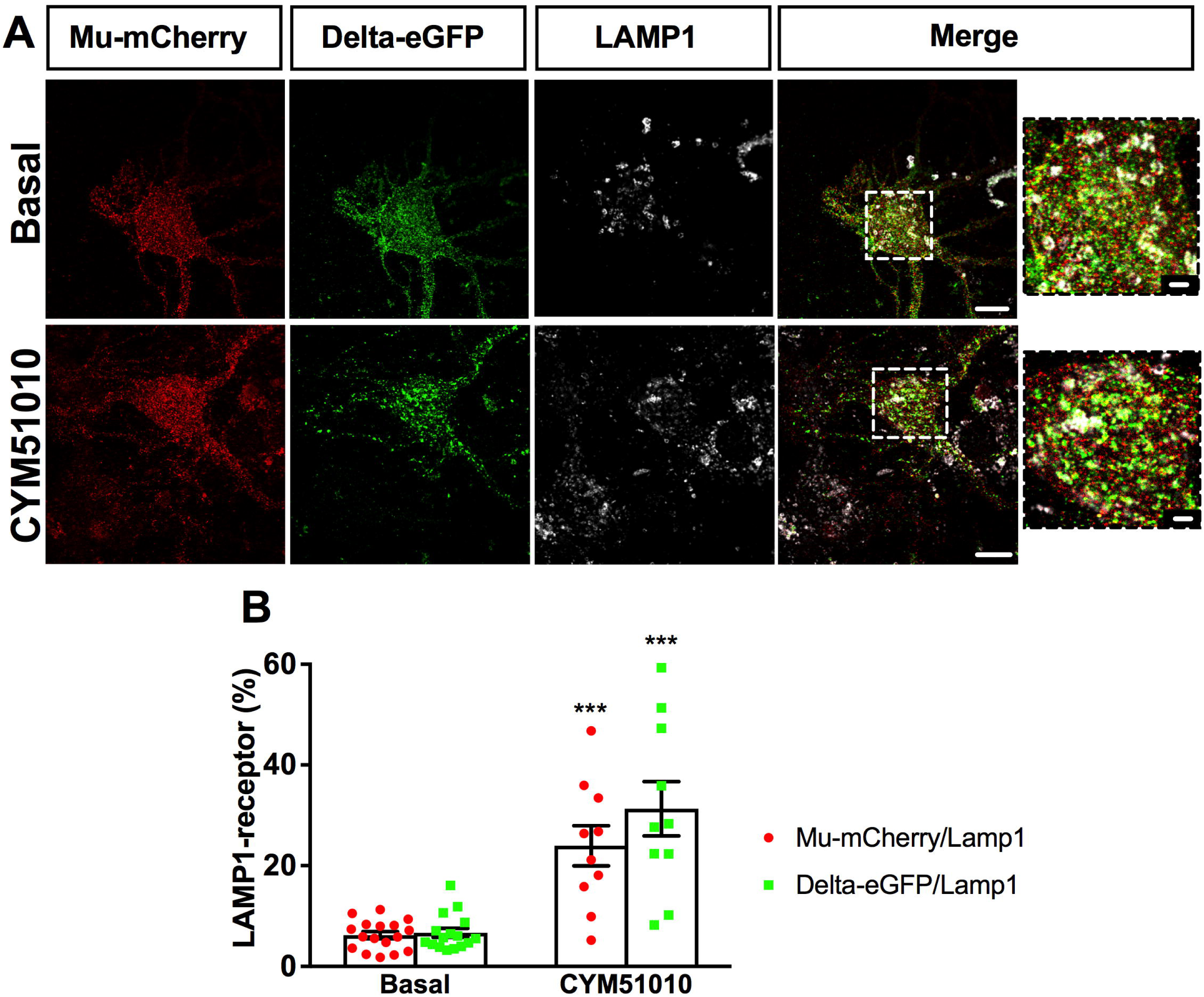
Mu and delta opioid receptors co-localize in the lysosomal compartment upon CYM51010 activation in primary hippocampal cultures. **A)** Representative confocal images showing mu-mCherry-delta-eGFP colocalization with LAMP1 immunoreactive compartment under basal condition or 60 minutes after CYM51010 application (400nM). Scale bar = 10µm (inset scale bar = 2.5 µm). **B)** Drug treatment induces statistically significant increase in the amount of colocalization of mu-mCherry / delta-eGFP colocalization with LAMP1 labeling. Two-way ANOVA F_drug treatment_ (1; 49)=62.70 ; p < 0.0001. F_receptor_ (1; 49) = 2.12, p = 0.15 ; F_interaction_ (1; 49)=3.65, p = 0.2. Tukey’s post hoc test: ***p < 0.001 for both mu-mCherry and delta-eGFP. N=10 to 20 neurons per group from at least 3 independent cultures.

We then sought to investigate whether internalization of the mu opioid receptor by CYM51010 was promoted by its association with the delta opioid receptor. In primary hippocampal cultures from single fluorescent knock-in animals expressing mu-mCherry and deficient for the delta opioid receptor, CYM51010 concentrations up to 1µM failed to induce mu-mCherry internalization (**Figure 3A-B**) with only limited mu opioid receptor clustering and subcellular redistribution at 10µM (**Figure 3B**). In contrast, the mu selective agonist DAMGO 1µM induced strong internalization of mu-mCherry **(see also Figure S2)** confirming the ability of this ligand to induce endogenous mu opioid receptor internalization (Trafton et al. 2000, Erbs et al. 2015).

**Figure 3:**
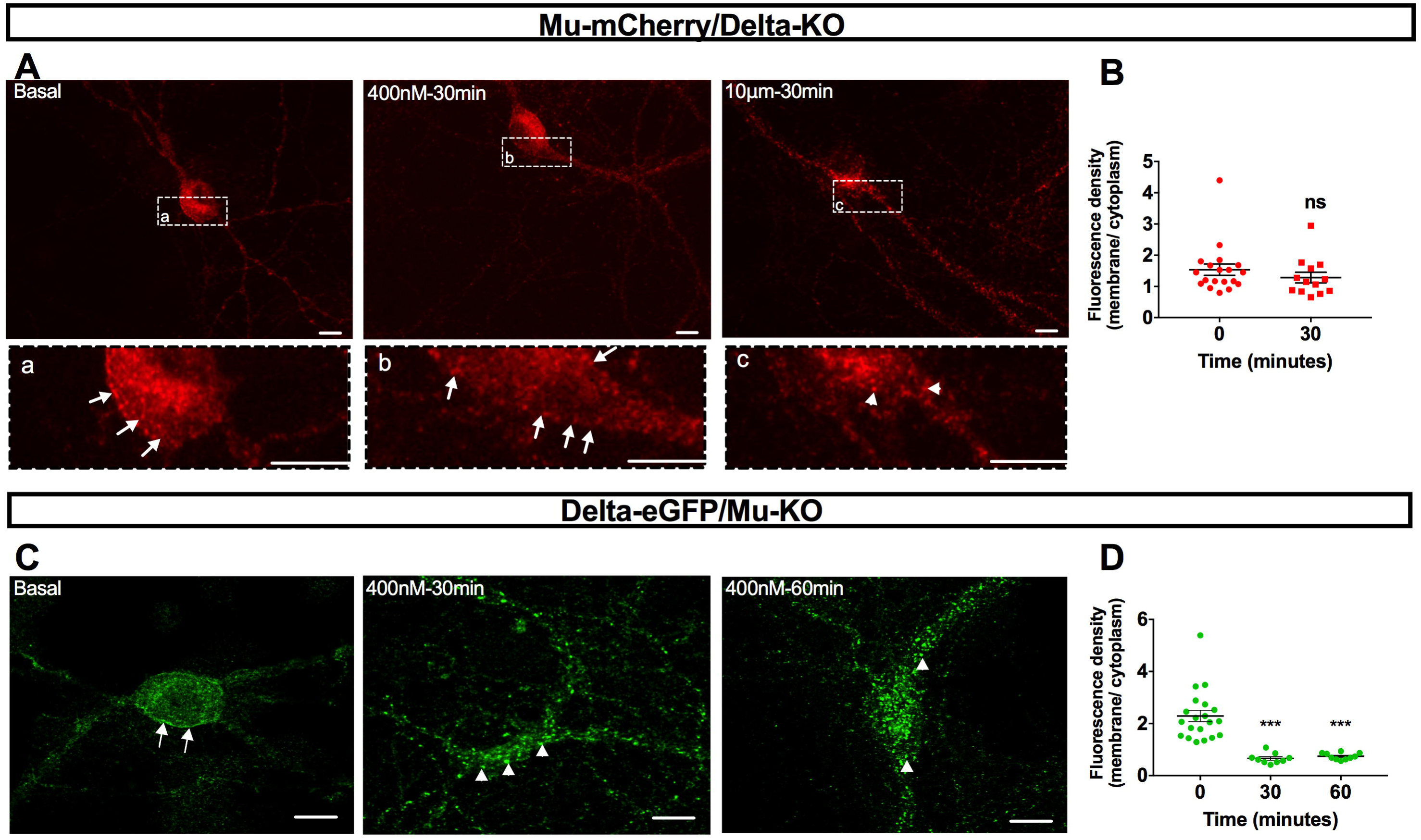
CYM51010 internalization of mu or delta opioid receptors in primary hippocampal cultures from mice deficient for one of the receptors. **A)** Representative confocal images showing that mu-mCherry is associated with the plasma membrane (arrows) in basal conditions and 30 minutes after CYM51010 (400nM) addition in delta-KO mice. Scale bar = 10µm **B)** Mu-mCherry internalization induced by CYM51010 application expressed as a ratio of membrane-associated versus intracellular fluorescence densities. Mann-Whitney test, p =0.20. N=13 to 20 neurons per group from at least 3 independent cultures. **C)** Representative confocal images showing that delta-eGFP is predominantly associated with the plasma membrane in basal conditions (arrows) in mu-KO mice whereas the association is mostly intracellular at 30 and 60 minutes after CYM51010 (400nM) addition (arrowheads). Scale bar = 10µm **D)** Delta-eGFP internalization induced by CYM51010 application expressed as a ratio of membrane-associated versus intracellular fluorescence densities. Kruskal-Wallis test (p <0.0001) followed by Dunn’s multiple comparison test Significant differences after multiple comparisons tests are expressed as p <0.001(***) compared to basal group. N=9-20 neurons per group from at least 3 independent cultures.

We then examined whether internalization of the delta opioid receptor upon activation by CYM51010 required mu opioid receptor co-expression. In primary hippocampal cultures from single fluorescent knock-in animals expressing delta-eGFP and deficient for the mu opioid receptor, CYM51010 (400nM) induced internalization of the delta opioid receptor (**Figure 3C**). In addition, predominant intracellular localization was observed 30 and 60 minutes after agonist application (**Figure 3D**) in agreement with kinetics described for the delta selective agonist SNC80 (Scherrer et al. 2006, Pradhan et al. 2009, Faget et al. 2012)) indicating that CYM51010 was able to promote delta opioid receptor internalization despite the lack of mu opioid receptor expression.

Together, this data establishes that mu opioid receptor internalization by CYM51010 is dependent on mu-delta receptor co-expression and directs the mu opioid receptor to the late endocytic compartment.

### CYM51010 induced mu-delta receptor co-internalization is blocked by pretreatment with mu or delta selective antagonists

In neurons expressing one receptor only, CYM51010 activation led to the internalization of delta but not mu opioid receptors (**Figure 3**). We therefore sought to determine whether co-internalization by CYM51010 required the two receptors to be in an active conformation. To this aim, we examined the impact of pre-treatment for 15 minutes with the mu selective antagonists β-FNA (20nM) or CTAP (200nM). Both antagonists prevented mu opioid receptor cellular redistribution but did not block delta opioid receptor internalization (**Figure 4A, C-D**). These results suggest that an active conformation of the mu opioid receptor is required for mu-delta co-internalization.

**Figure 4:**
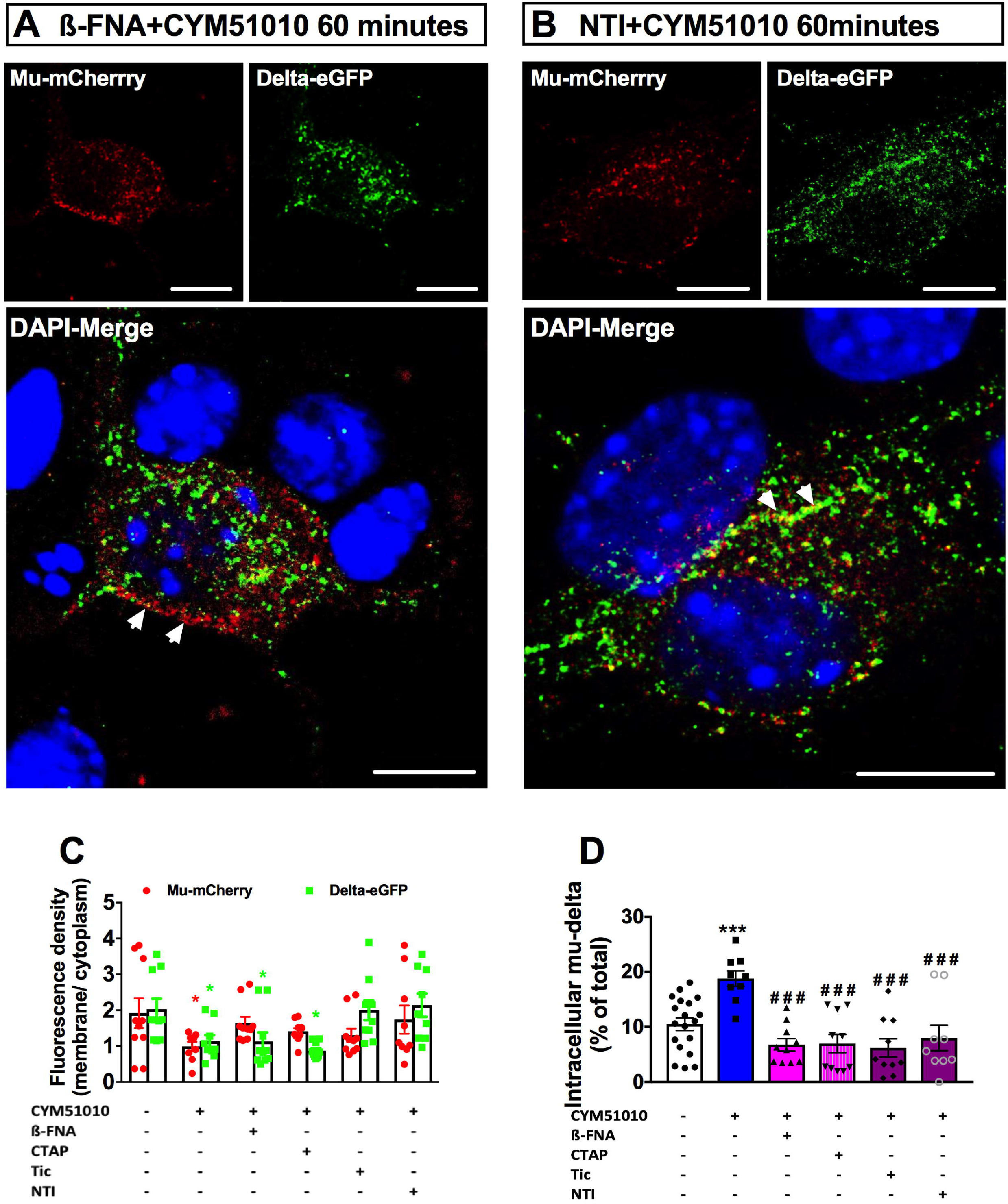
Antagonist pre-treatment abolishes mu-delta opioid receptor co-internalization by CYM51010 in primary hippocampal cultures. **A)** Representative confocal images showing mu-mCherry predominant localization at the plasma membrane and delta-eGFP extensive internalization after pre-treatment with the mu antagonist β-FNA (200nM) for 15 minutes followed by incubation for 60 minutes with CYM51010 (400nM). Scale bar = 10µm. **B)** Representative confocal images showing mu-mCherry and delta-eGFP predominant localization at the plasma membrane after pre-treatment with delta antagonist naltrindole (200nM) (NTI) for 15minutes followed by incubation for 60 minutes with CYM51010 (400nM). Scale bar = 10µm. **C)** Pre-treatment with the mu antagonists β-FNA or CTAP (200nM) blocks mu-mCherry but not delta-eGFP internalization, whereas pre-treatment with the delta antagonists naltrindole (NTI) and Tic-deltorphin (tic) (200nM) prevent internalization of both mu-mCherry and delta-eGFP. Receptor internalization is expressed as a ratio of membrane-associated versus intracellular fluorescence densities for each receptor. Two-way ANOVA F_treatment_ (5;104)=4.73 ; p =0.0001. F_receptor_ (1; 104)=0.1; p= 0.84 ; F_interaction_ (5; 100)=1.96; p = 0.0006. Multiple comparisons with Tukey’s posthoc test *p=0.04 basal vs CYM51010 for mu-mCherry, *p=0.04 basal vs CYM51010, *p =0.04, basal vs β-FNA, *p = 0.04, basal vs CTAP, **p = 0.006. for delta-eGFP. N=9 to 20 neurons per group from at least 3 independent cultures. **D)** Mu-mCherry/delta-eGFP co-internalization is prevented by treatment with either mu or delta antagonists. Percentage of colocalized receptors in the cytoplasm after drug treatment. The fraction of cytoplasmic mu-delta heteromers is expressed as the percentage of mu-mCherry and delta-eGFP overlapping objects detected in vesicle-like structures 60 minutes after CYM51010 application. One-way ANOVA (p<0.0001) followed by multiple comparisons Dunnett’s test. Significant differences after multiple comparisons tests are expressed as ***p<0.001 when compared to basal group and ^###^p<0.001 when compared to CYM51010 without antagonists. N=9 to 20 neurons per group from at least 3 independent cultures.

We then evaluated the need for delta opioid receptor activation in the co-internalization process. Whereas pre-treatment with mu antagonists blocked mu but not delta opioid receptor internalization, pre-treatment with the selective delta antagonists naltrindole or Tic-deltorphin 200 nM blocked the internalization of both delta and mu opioid receptors (**Figure 4B-D**). This indicates that mu-delta cellular redistribution is driven by delta opioid receptor expression and activation. Together, this data indicates that mu-delta receptor co-internalization upon CYM 51010 activation is driven by delta opioid receptors and requires both mu and delta opioid receptors to be in an active conformation.

### CYM51010 induces phosphorylation of mu and delta opioid receptors

To confirm that active conformation of the receptors is required for internalization, we also examined the phosphorylation profile of mu and delta opioid receptors *in vivo* in response to CYM51010 (10mg/kg i.p.) using antibodies for the different phosphorylation sites on the receptors. The mu-delta agonist induced phosphorylation of the serine residue S363, the primary site of the delta opioid receptor whether the receptor was expressed alone (delta-eGFP/mu KO mouse) or together with the mu opioid receptor (mu-mCherry/ delta-eGFP mouse) (**Figure 5A**). This result is similar to the phosphorylation following activation by the delta selective agonist SNC80 (**Figure 5A**). CYM51010 also promoted phosphorylation at the primary mu opioid receptor phosphorylation site serine S375 and at threonine residues T370, T376, T379 whether the mu opioid receptor was expressed alone (mu-mCherry/delta KO mouse) or together with the delta opioid receptor (mu-mCherry/ delta-eGFP mouse). The phosphorylation profile was similar to the one observed following activation with the mu selective agonist fentanyl (0.3mg/kg i.p.) (**Figure 5B**).

**Figure 5:**
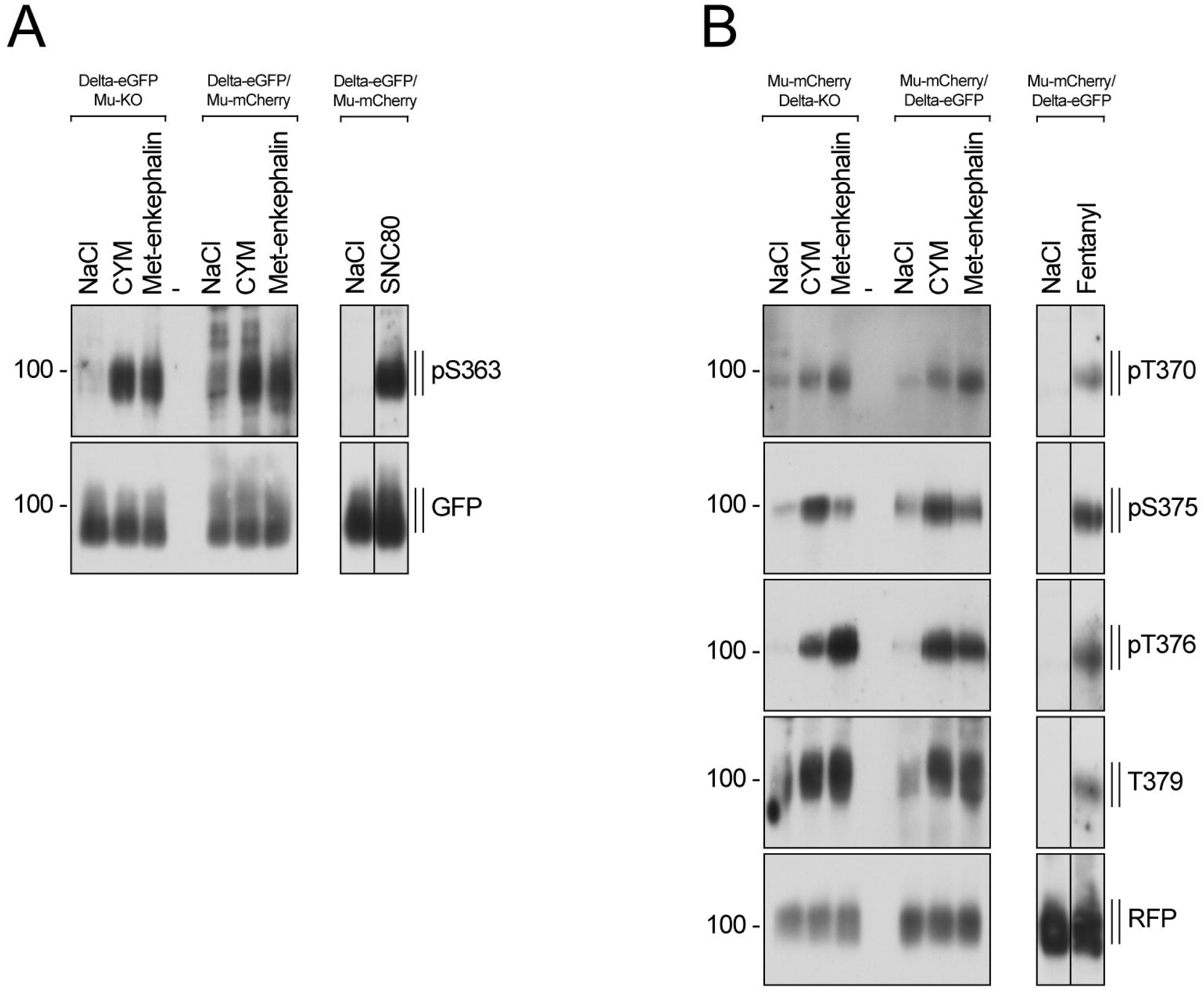
Agonist-induced *in vivo* phosphorylation of mu and delta opioid receptors. **A)** Representative Western blot showing the phosphorylation of the delta opioid receptor at the C-terminal serine 363 residue upon by CYM51010 (10mg/kg i.p,), SNC80 (10mg/kg i.p.) or Met-enkephalin (375 nmol i.c.v.) *in vivo* administration in double fluorescent knock-in mice (delta-eGFP/mu-mCherry) or in mice only expressing the fluorescent delta opioid receptor (delta-eGFP/mu KO). **B)** Representative Western blot showing the phosphorylation of the mu opioid receptor at specific C-terminal residues upon CYM51010 (10mg/kg i.p,), fentanyl (0.3mg/kg i.p.) or Met-enkephalin (375 nmol i.c.v.) *in vivo* administration in double fluorescent knock-in mice (delta-eGFP/mu-mCherry) or in mice only expressing the fluorescent mu opioid receptor (mumCherry/delta KO).

Together, this data indicates that mu-delta receptor co-internalization upon CYM 51010 requires phosphorylation of both mu and delta opioid receptors.

### Mu-delta receptor co-internalization is ligand specific

We then examined whether other synthetic opioid agonists were able to promote mu-delta receptor co-internalization in primary hippocampal cultures. Mu-delta receptor co-localization in the cytoplasm was increased 30 minutes after stimulation with the mu agonist DAMGO (1µM) or the delta agonist deltorphin II (100nM) but not upon stimulation with the delta agonist SNC 80 (**Figure 6**). These data establish ligand specific internalization of endogenous mu-delta heteromers by exogenous opioids.

**Figure 6:**
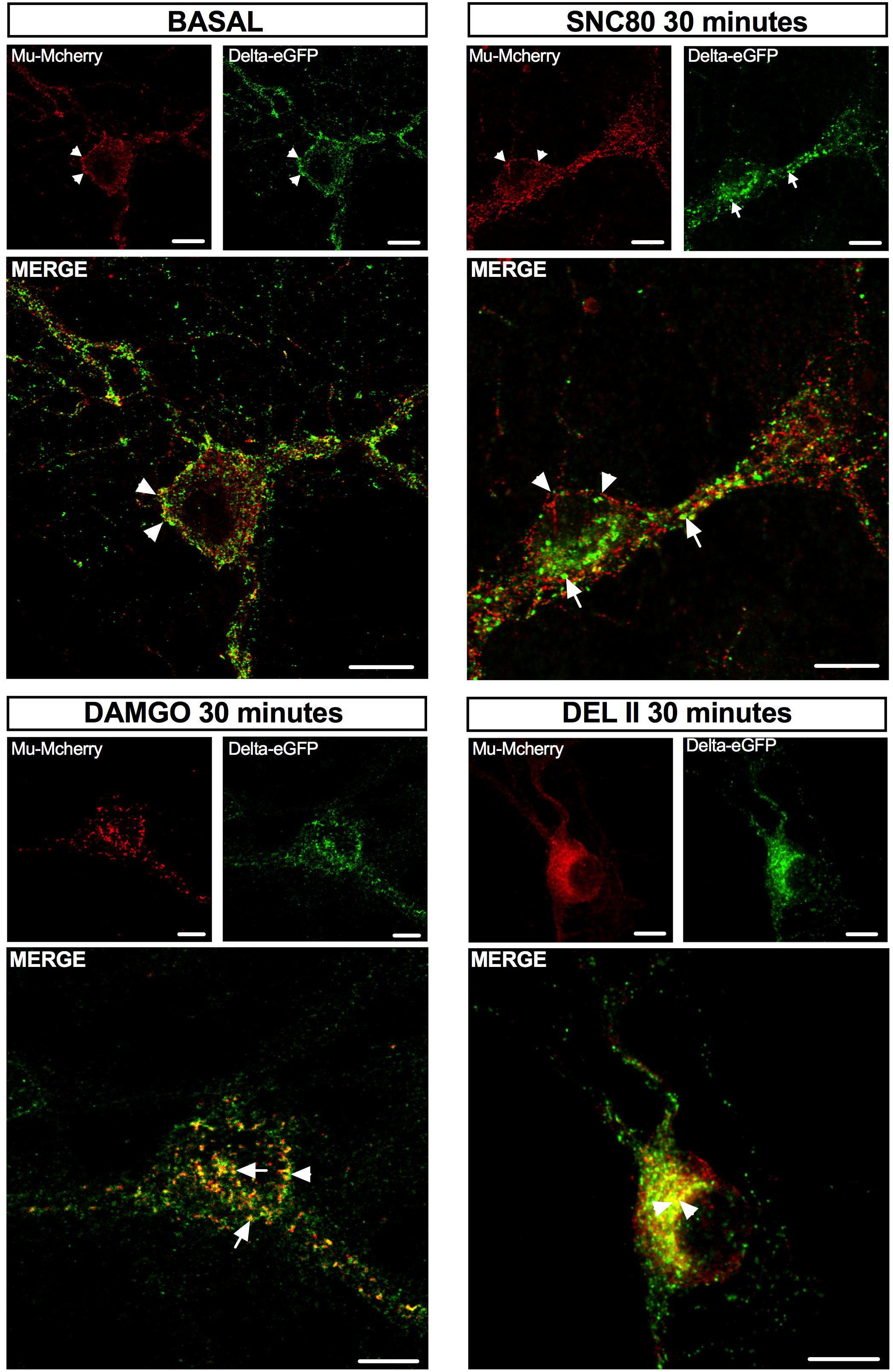
Mu-delta opioid receptor co-internalization is ligand selective in primary hippocampal cultures. Representative confocal images showing mu-mCherry and delta-eGFP fluorescence at the plasma membrane (arrowheads) under basal conditions or co-internalized in vesicle-like structures (arrows) 30 minutes after DAMGO (1 µM) or deltorphin II (100nM) but not SNC80 (100nM). Scale bar =10µm.

### Met-enkephalin induced mu-delta co-internalization and co-localization in the late endosomal compartment

The endogenous opioid peptide met-enkephalin promotes internalization and fast recycling to the plasma membrane of mu opioid receptors (Trafton et al. 2000, Minnis et al. 2003) as well as internalization and subsequent lysosomal degradation of the delta opioid receptor (Scherrer et al. 2006, Pradhan et al. 2009). Accordingly, in primary hippocampal cultures from single fluorescent knock-in mice expressing mu-mCherry and deficient for the delta opioid receptor, activation with met-enkephalin 100nM led to mu-mCherry predominant intracellular localization 30 minutes after agonist application. After 60 minutes, mu-mCherry was detected at the plasma membrane consistent with the kinetics of a fast recycling receptor (**Figure 7A, B**). In contrast, in primary hippocampal cultures from single fluorescent knock-in mice expressing delta-eGFP and deficient for the mu opioid receptor, the receptor was mainly located in the cytoplasm at both time points following activation with met-enkephalin 100nM consistent with the kinetics of a receptor type that is degraded upon internalization (**Figure 7C, D**).

**Figure 7:**
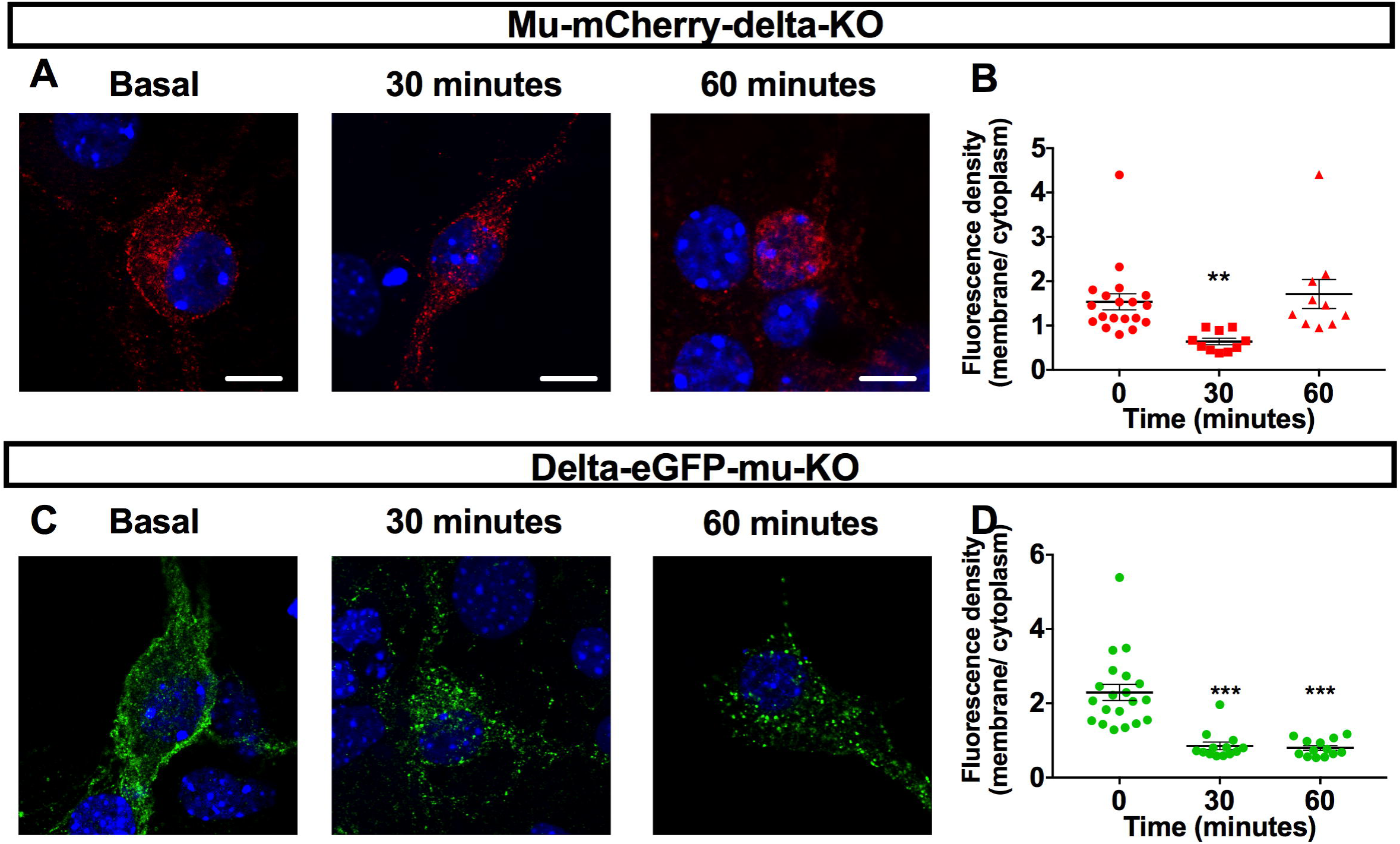
Met-enkephalin internalization of mu or delta opioid receptors expressed alone. **A)** Representative confocal images showing mu-mCherry fluorescence mainly localized at the plasma membrane under basal conditions or internalized in vesicle-like structures 30 minutes after Met-enkephalin challenge (100nM). 60 minutes, mu-mCherry was mainly detected back at the plasma membrane. Scale bar = 10µm. **B)** Quantification of mu-mCherry internalization expressed as a ratio of membrane-associated versus intracellular fluorescence densities. Kruskal-Wallis test (p <0.0001) followed by Dunn’s multiple comparison test p<0.001(***) compared to basal group. n=10 to 19 neurons per group from at least 3 independent cultures. **C)** Representative confocal images showing delta-eGFP fluorescence mainly localized at the plasma membrane under basal conditions or internalized in vesicle-like structures 30 or 60 minutes after Met-enkephalin challenge (100nM). Scale bar = 10µm. **D)** Quantification of delta-eGFP internalization expressed as a ratio of membrane-associated versus intracellular fluorescence densities. Kruskal-Wallis test (p <0.0001) followed by Dunn’s multiple comparison test. p=0.002(**) compared to basal group. n=13 to 20 neurons per group from at least 3 independent cultures.

We then examined whether met-enkephalin also promoted mu-delta receptor co-internalization and modified the mu opioid receptor intracellular fate. Met-enkephalin (100nM) application for 15, 30 or 60 minutes was associated with reduced fluorescence at the cell surface and led to the redistribution of both mu-mCherry and delta-eGFP signals in cytoplasmic vesicular structures (**Figure 8A, B**). Importantly, the percentage of mu-delta heteromers also decreased at the cell surface and increased intracellularly at the three time points (**Figure 8C)** with significant increase in mu-delta heteromers at 60 minutes **(Fig. 8D**). At this time point, both mu and delta opioid receptors co-localized with the late endosomallysosomal marker LAMP1 (**Figure 9**).

**Figure 8:**
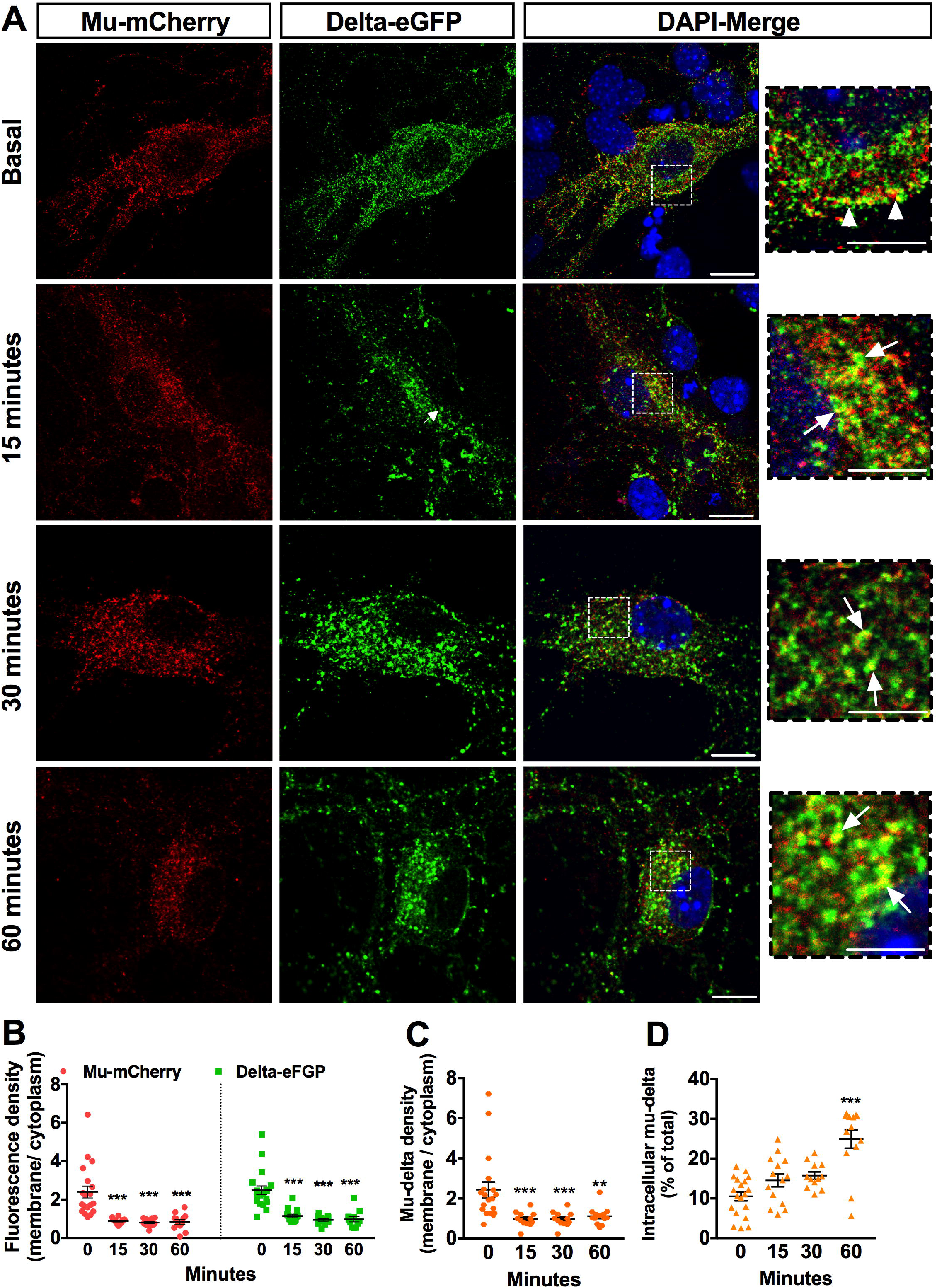
Mu and delta opioid receptors co-internalize upon met-enkephalin activation in primary hippocampal cultures. **A)** Representative confocal images showing mu-mCherry and delta-eGFP subcellular localization. In basal conditions, receptors are predominantly co-localized (inset arrows) at the periphery of the neuron (top panel). Following 100nM Met-enkephalin application for 15, 30, 60 minutes, receptors are co-internalized and co-localize in the cytoplasm (inset arrows). Scale bar = 10 µm (inset scale bar = 5 µm). **B)** Receptor internalization induced by met-enkephalin application (100nM) expressed as a ratio of membrane-associated versus intracellular fluorescence densities for each receptor. Two-way ANOVA F_drug treatment_ (3;105)=34.54 ; p <0.0001. F_receptor_ (1; 105)=1.27 ; F_interaction_ (3; 105)=0.08. Multiple comparisons were made with Tukey’s test, ***: p <0.001. N=12 to 19 neurons per group from at least 3 independent cultures. **C)** Subcellular redistribution of mu-delta heteromers expressed as a ratio of membrane-associated versus intracellular fluorescence densities for co-localized mu-mCherry and deltaeGFP receptors. Kruskal Wallis test (p < 0.0001) followed with multiple comparisons Dunn’s test. ** p <0.01 ; ***p <0.001. N=12 to 19 neurons per group from at least 3 independent cultures. **D)** Fraction of cytoplasmic mu-delta heteromers expressed as the percentage of mumCherry and delta-eGFP overlapping objects detected in vesicle-like structures at the different times. Kruskal Wallis test (p < 0.0001) followed with multiple comparisons Dunn’s test. p = 0.28 15 min vs basal, p = 0.09, 30 min vs basal, ***p < 0.0001 60 min vs basal. N=12 to 19 neurons per group from at least 3 independent cultures.

**Figure 9:**
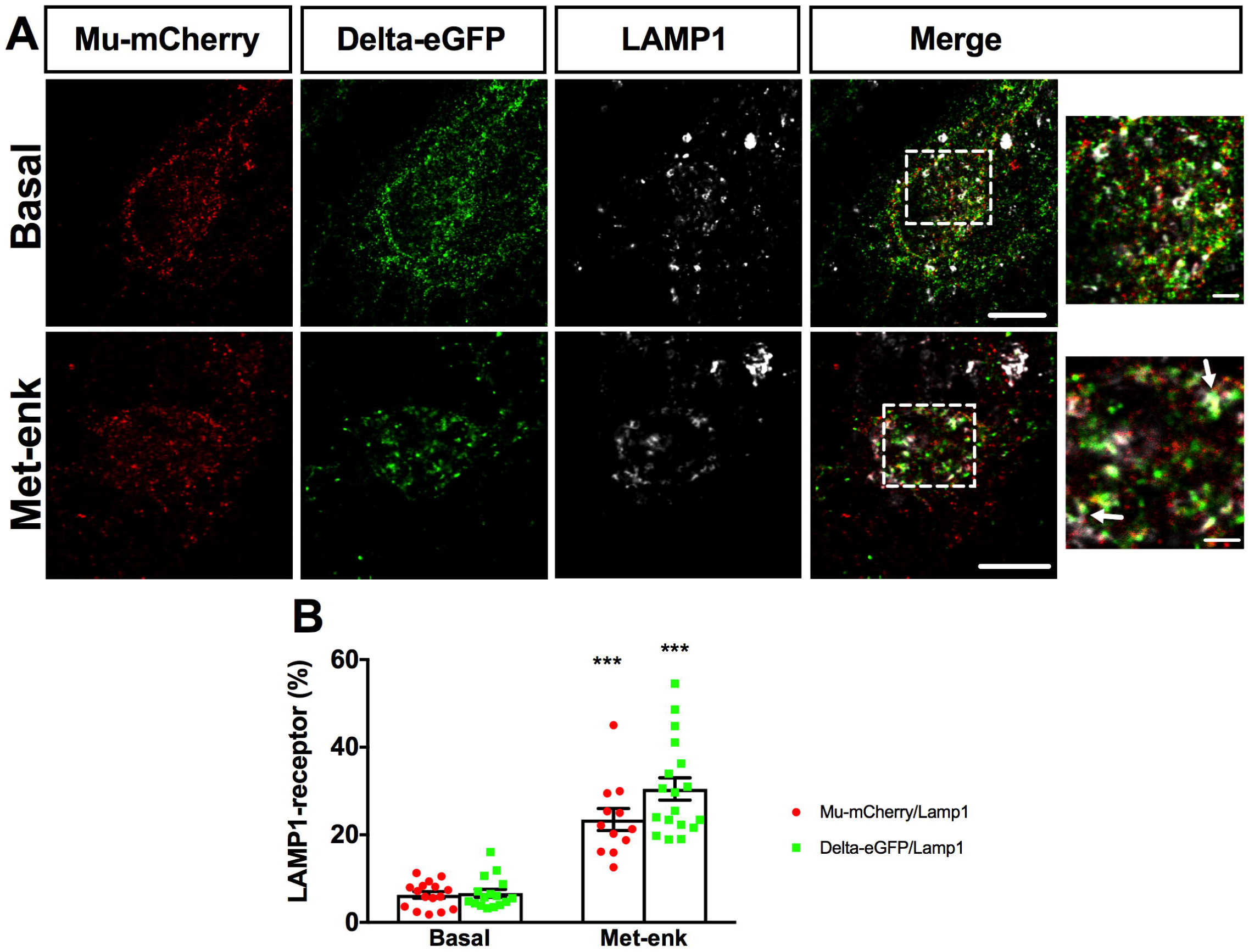
Mu and delta opioid receptors co-localize in the lysosomal compartment upon met-enkephalin activation in primary hippocampal cultures. **A)** Representative confocal images showing mu-mCherry-delta-eGFP colocalization with LAMP1 immunoreactive compartments under basal condition or 60 minutes after Metenkephalin (100nM) application. Scale bar = 10 µm (Inset Scale bar = 2.5 µm). **B)** Met-enkephalin induces statistically significant increases in the amount of mu-mCherry / delta-eGFP colocalization with LAMP1 labeling. Two-way ANOVA F_drug treatment_ (1;59)=123.5 ; P < 0.0001. F_receptor_ (1; 59) = 4.04, p-value = 0.05 ; F_interaction_ (1; 59)=3.08, p-value = 0.08. Tukey’s multiple comparisons: ***p < 0.0001 for both mu-mCherry and delta-eGFP. N=12 to 16 neurons per group from at least 3 independent cultures.

Our results therefore suggest that mu and delta opioid receptors are targeted together to the degradation pathway as observed upon activation with CYM51010. They also establish that the endogenous opioid peptide met-enkephalin shifts the mu opioid receptor intracellular fate from the fast recycling to the degradation mode upon mu-delta heteromerization.

### Met-enkephalin induced mu-delta receptor co-internalization is blocked by pretreatment with mu or delta selective antagonists

We then examined the impact of mu or delta selective antagonists on mu-delta receptor co-internalization in response to met-enkephalin. A 15 minutes pre-treatment with the mu selective antagonists CTAP or β-FNA prevented changes in the mu opioid receptor subcellular distribution with mu-mCherry fluorescence predominantly remaining visible at the plasma membrane **(Figure 10A)**. In contrast, delta-eGFP was internalized and detected in intracellular vesicles after 60 minutes **(Figure 10B)**. Pre-treatment with the mu antagonists thus prevented co-internalization indicating that the mu opioid receptor needs to be in an active form to be co-internalized with the delta opioid receptor (**Figure 10C, D**). Pretreatment with the delta selective antagonists naltrindole or Tic-deltorphin prevented delta opioid receptor internalization but also blocked mu opioid receptor subcellular redistribution indicating that the delta opioid receptor acted as the driver of the co-internalization (**Figure 10B-D**). Similarly, pre-treatment with pertussis toxin (100ng/ml) for 24 hours, known to promote receptor uncoupling from inhibitory Gi/o proteins through ADP ribosylation of the Gα subunit, also prevented mu-delta co-internalization **(see Figure S3)**.

**Figure 10:**
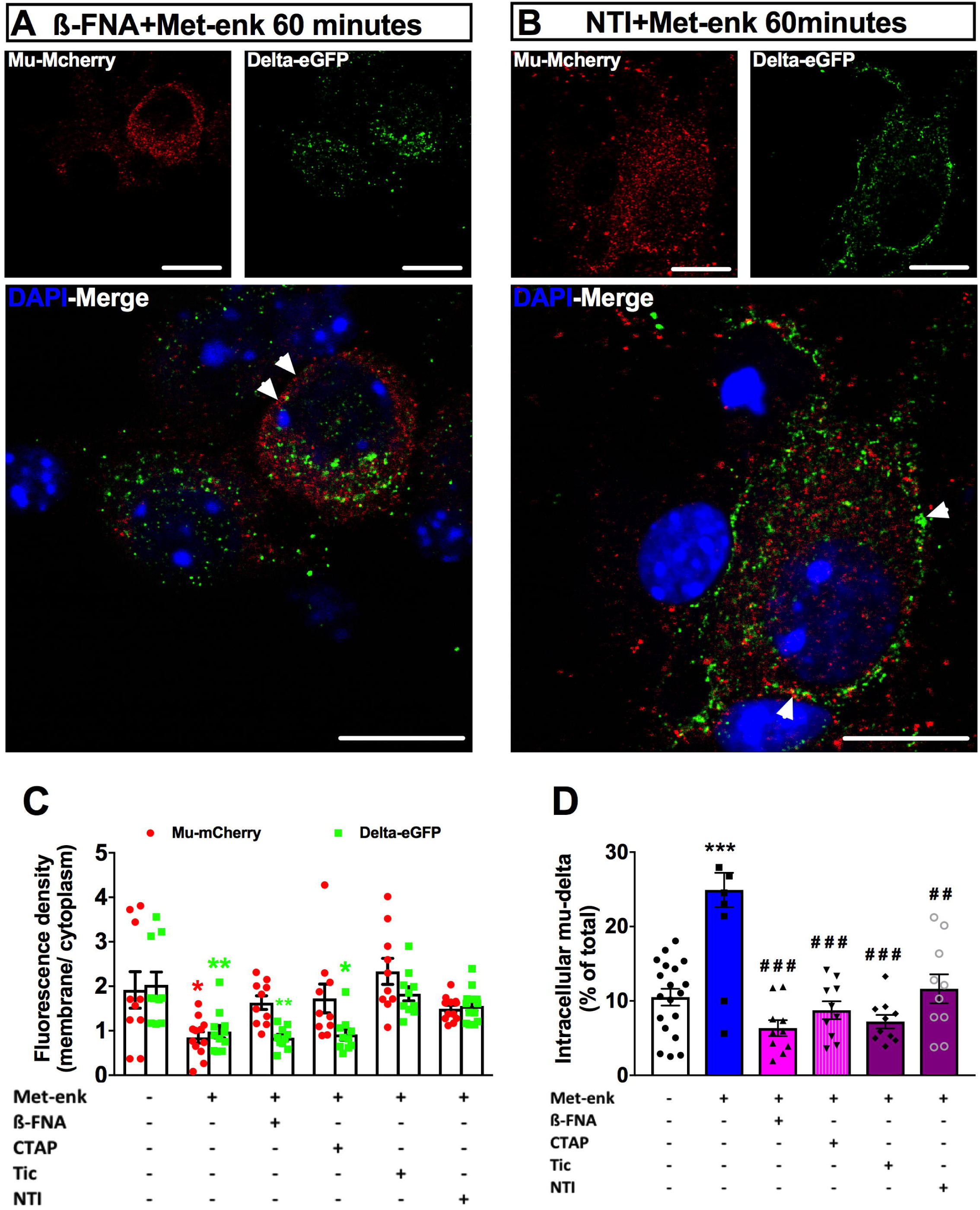
Antagonist pre-treatment abolishes mu-delta opioid receptor co-internalization by met-enkephalin in primary hippocampal cultures. **A)** Representative confocal images showing mu-mCherry predominant localization at the membrane and delta-eGFP extensive internalization following met-enkephalin (100nM) incubation for 60 minutes in primary neurons pretreated with the mu antagonist β-FNA 200 nM for 15 minutes. Scale bar = 10µm. **B)** Representative confocal images showing mu-mCherry and delta-eGFP predominant localization at the membrane following met-enkephalin incubation for 60 minutes in primary neurons pretreated with the delta antagonist naltrindole (NTI) (200nM) for 15 minutes Scale bar = 10µm. **C)** Quantification of mu or delta opioid receptor internalization respectively, expressed as a ratio of receptor density in the membrane versus mu or delta mu-mCherry or delta-eGFP receptor density in the cytoplasm. Pre-treatment with the mu selective antagonists β-FNA (20 nM) or CTAP (200nM) blocks mu-mCherry but not delta-eGFP internalization following met-enkephalin application. Pre-treatment with the delta selective antagonists NTI or Tic-deltorphin (200nM) blocks met-enkephalin-induced internalization of both mu-mCherry and delta-eGFP. Two-way ANOVA F_drug treatment_ (5;100)=6.38 ; p <0.0001. F_receptor_ (1; 100)=3.96; p= 0.04 ; F_interaction_ (5; 100)=1.97; p = 0.09. N=10 to 12 neurons per group from at least 3 independent experiments. **D)** Mu-delta co-internalization is prevented by treatment with either mu or delta selective antagonists. Percentage of co-localized receptors in the cytoplasm after drug treatment. The fraction of cytoplasmic mu-delta heteromers is expressed as the percentage of mu-mCherry and delta-eGFP overlapping objects detected in vesicle-like structures 60 minutes after CYM51010 application. Kruskal Wallis test (p = 0.0001) followed by multiple comparisons Dunn’s test. Significant differences after multiple comparisons tests are expressed as ***, p < 0.001when compared to basal group and ###, p < 0.001 when compared to Met-enkephalin without antagonists. N=10 to 12 neurons per group from at least 3 independent cultures.

Data indicates that mu-delta co-internalization following met-enkephalin activation involves active conformations of both receptors similar to that produced by CYM51010.

### Met-enkephalin induces mu-delta receptor co-internalization and co-localization in the late endosomal compartment in vivo

We then examined the *in vivo* intracellular fate of mu-delta heteromers in response to met-enkephalin administration (375 nmoles i.c.v.). Under basal conditions, mu and delta opioid receptors were predominantly located at the plasma membrane (**Figure 11A, B**) with about 15% of mu and delta opioid receptors co-localized (**Figure 11C**), suggesting mu-delta heteromerization at the neuronal surface. These values were very similar to those collected in primary hippocampal cultures (**Figure 1, see also Figure S1**). Met-enkephalin administration led to the sequestration of both mu and delta opioid receptors (**Figure 11A, B**) and maintained the extent of receptor co-localization in the cytoplasm (**Figure 11C**). In addition, mu and delta opioid receptors co-localized with the late endosomal-lysosomal LAMP1 marker 60 minutes after met-enkephalin administration, indicative of mu-delta receptor co-internalization **(Figure 11D)**.

**Figure 11:**
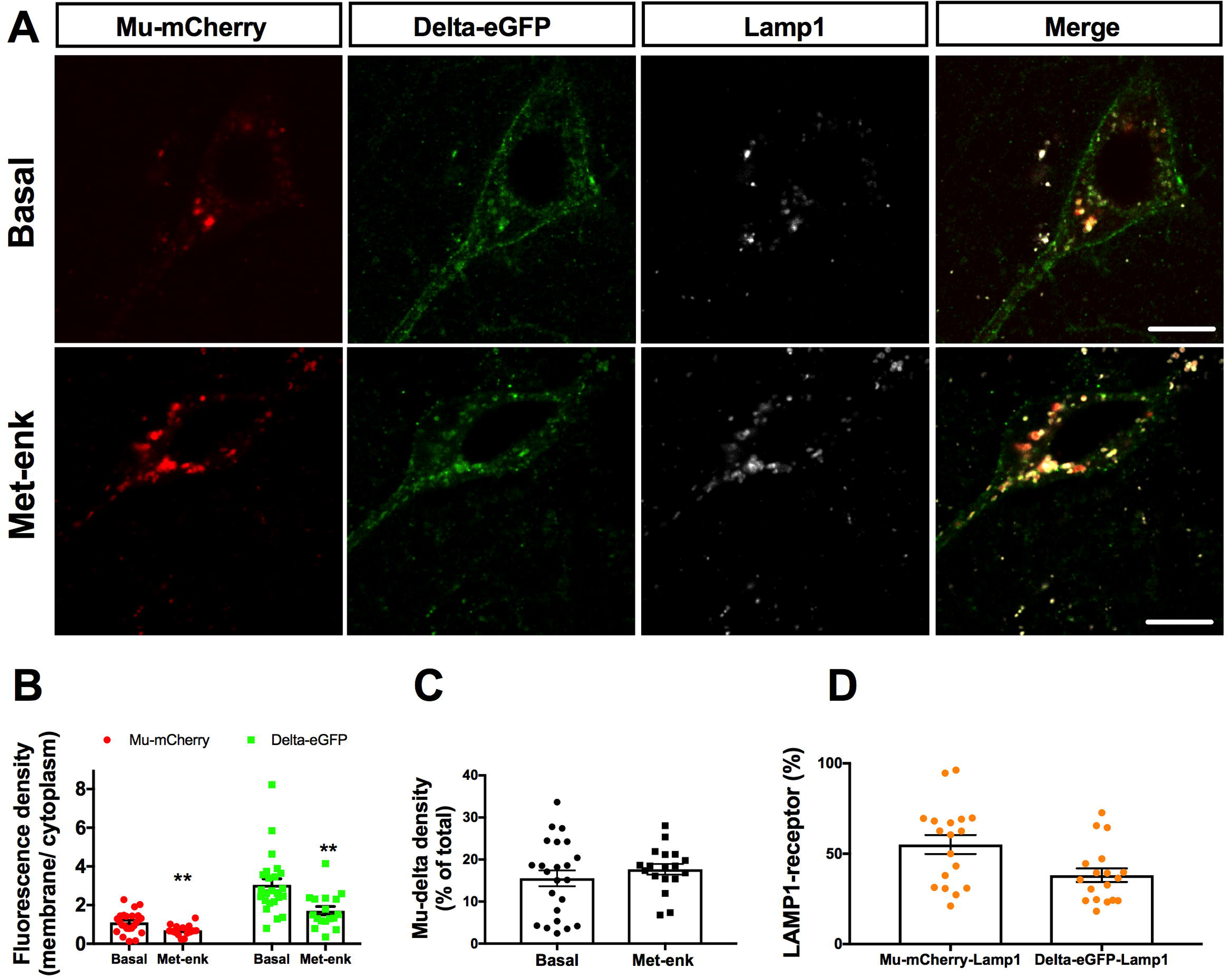
Mu and delta opioid receptors co-localize in the lysosomal compartment upon Met-enkephalin activation *in vivo*. **A)** Representative confocal images showing mu-mCherry and delta-eGFP co-localization with LAMP1 immunoreactive compartment in the hippocampus under basal condition or 60 minutes after Met-enkephalin (100nM) application. Scale bar = 10µm. **B)** Receptor internalization induced by Met-enkephalin injection (375 nmoles i.c.v.) expressed as a ratio of membrane-associated versus intracellular fluorescence densities for each receptor. Student’s unpaired t-test, **p = 0.003 for mu-mCherry, ***p = 0.001 for delta-eGFP. N=17 to 25 neurons per group from at least 3 mice. **C)** Mu-delta heteromers expressed as the percentage of mu-mCherry and delta-eGFP overlapping objects detected at the plasma membrane or in the cytoplasm in basal conditions Student’s unpaired t-test test (p > 0.05) N=17 to 25 neurons per group from at least 3 mice. **D)** Met-enkephalin increases the amount of mu-mCherry / delta-eGFP co-localization with LAMP1 labeling after 60 minutes. Student’s unpaired t-test test. N=18 neurons per group from at least 3 mice.

Endogenous mu-delta heteromers are thus present at the plasma membrane *in vivo* and undergo similar intracellular fate in the hippocampal tissue or in primary hippocampal cultures.

### Met-enkephalin induces phosphorylation of mu and delta opioid receptors

To confirm activation of either receptor, we examined their phosphorylation profile following met-enkephalin activation *in vivo* (375 nmoles i.c.v.). Strong phosphorylation of the serine residue S363 was observed whether the receptor was expressed alone (delta-eGFP/mu KO mouse) or together with the mu opioid receptor (mu-mCherry/ delta-eGFP mouse) **(Figure 5A)**. Similarly, robust phosphorylation at residues S375, T370, T376 and T379 was observed whether the mu opioid receptor was expressed alone (mu-mCherry/delta KO mouse) or together with the delta opioid receptor (mu-mCherry/ delta-eGFP mouse) **(Figure 5B)**.

Together, data indicates that internalization of mu-delta heteromers following metenkephalin activation requires delta opioid receptors and involves active conformations of both receptors similar to that produced by CYM51010. Together, this data indicates that mu-delta receptor co-internalization induced by met-enkephalin requires phosphorylation of both mu and delta opioid receptors as shown for co-internalization induced by the mu-delta biased agonist CYM51010.

### β-endorphin does not promote mu-delta opioid receptor co-internalization

We then investigated the internalization profile of mu and delta opioid receptors upon activation by β-endorphin in primary hippocampal cultures. We observed a strong decrease of the delta-eGFP signal at the plasma membrane that persisted for at least 60 minutes after β-endorphin (100nM) application in agreement with the slow recycling mode of the delta opioid receptor (**Figure 12A, B**). In contrast, changes in the mu-mCherry fluorescence associated with the plasma membrane were transient with a strong decrease in the signal after 15 minutes followed by partial recovery at 30 and 60 minutes as expected for fast receptor recycling to the plasma membrane (**Figure 12A, B**). Clear segregation of mu-mCherry and delta-eGFP signals was observed (**Figure 12A**). The density of mu–delta heteromer at the plasma membrane was not statistically significantly decreased at the 15 minute time point (**Figure 12C**) and the density of intracellular mu-delta heteromers did not increase over time (**Figure 12D**).

**Figure 12:**
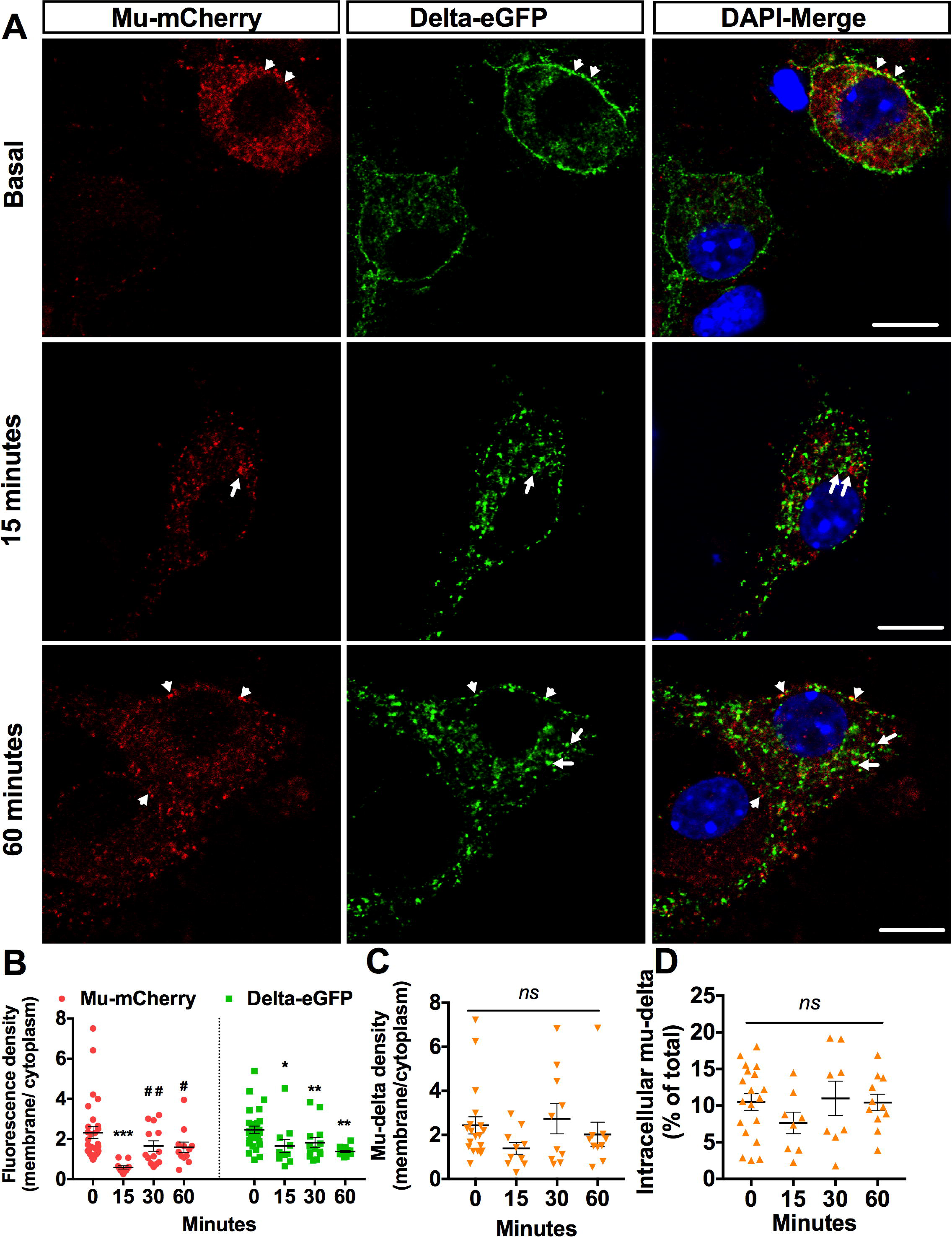
Mu and delta opioid receptors do not co-internalize upon β-endorphin activation in primary hippocampal cultures. **A)** Representative confocal images showing mu-mCherry and delta-eGFP subcellular localization. In basal conditions receptors are predominantly co-localized (arrowheads) at the surface of the neuron. Upon activation with β-endorphin (100nM), receptors are internalized (arrows) but no significant co-localization can be detected. Moreover, mu-mCherry is detectable at the plasma membrane after 60 minutes (arrowheads). Scale bar = 10µm. **B)** Quantification of internalization given as a ratio of mu-mCherry or delta-eGFP receptor density in the membrane relative to the density in the cytoplasm. Two-way ANOVA F _treatment_ (3; 123) = 9.61 ; p-value <0.0001. F_receptor_ (1; 123)=2.20, p-value= 0.19 ; F_interaction_ (3; 123)=0.55, p-value=0.20. Multiple comparisons were made with Tukey’s test. For Mu-mCherry: *** p < 0.001 15 min vs basal, p = 0.15 30 min vs basal; p = 0.11 60 min vs basal. ## p = 0.03 30 min vs 15 min; # p = 0.04 60 min vs 15 min. For Delta-eGFP: ***p = 0.001 15min vs basal, *p = 0.02 30 min vs basal, ***p = 0.001 60 min vs basal. N=10 to 20 neurons per group from at least 3 independent cultures. **C)** Subcellular redistribution of mu-delta heteromers expressed as a ratio of membrane-associated versus intracellular fluorescence densities for co-localized mu-delta opioid receptors. Kruskal Wallis test (p = 0.21) followed with multiple comparisons Dunn’s test. NS= No statistically significant differences (p = 0.27 15 min vs basal, p = 0.94 30 min vs basal, p = 0.86 60min vs basal). N=10 to 20 neurons per group from at least 3 independent experiments. **D)** Fraction of cytoplasmic mu-delta heteromers expressed as a percentage of mu-mCherry and delta-eGFP overlapping objects detected in vesicle-like structures at the different time points. Kruskal Wallis test (p = 0.51) followed with multiple comparisons Dunn’s test. NS= No statistically significant differences (p = 0.47 15 min vs basal, p = 0.99 30 min or 60 min vs basal). N=10 to 20 neurons per group from at least 3 independent cultures.

Together, this data establishes that stimulation with β-endorphin does not induce co-trafficking of mu and delta opioid receptors but rather each receptor maintains its respective fast and low recycling modes. Unlike met-enkephalin, β-endorphin does not alter the mu opioid receptor intracellular fate through mu-delta heteromer activation.

### The kinetics of Met-enkephalin induced ERK1/2 phosphorylation is altered by mu-delta heteromerization

Mu opioid agonists promote rapid and transient ERK1/2 phosphorylation via activation of the G protein dependent pathway (Massotte et al. 2002). Heteromerization of mu-delta opioid receptors in co-transfected CHO cells shifted ERK1/2 phosphorylation towards a later and more sustained profile following DAMGO stimulation (Rozenfeld and Devi 2007). Using the SKNSH cell line co-expressing endogenous mu and delta opioid receptors, the authors also showed that this profile was β-arrestin 2 dependent (Rozenfeld and Devi 2007). We therefore investigated whether mu-delta heteromerization also affected the kinetics of ERK1/2 phosphorylation in primary hippocampal neurons. As expected, DAMGO (1µM) induced robust increase in ERK1/2 phosphorylation that lasted for 10 minutes before slowly going back to basal levels after 20 minutes (**Figure 13A**). Activation by the mu-delta biased agonist CYM51010 (400nM) led to a rapid 50% increase in ERK1/2 phosphorylation that lasted for 15 minutes before returning to baseline (**Figure 13A**), consistent with its ability to recruit β-arrestins (Gomes et al. 2013). We then examined ERK1/2 phosphorylation following activation by endogenous opioid peptides. Activation by β-endorphin (100nM) was slower compared to the other ligands tested and led to 40% increase in ERK1/2 phosphorylation 5 minutes after stimulation before slowly returning to baseline levels at 20 minutes (**Figure 13A**). Remarkably, met-enkephalin (100nM) induced a biphasic phosphorylation kinetics with a rapid transient 50% increase in phosphorylation after 3 minutes followed by a stronger and sustained phase that persisted for up to 20 minutes (**Figure 13A)**. This data indicates that activation of mu-delta heteromer by all four ligands promoted ERK1/2 phosphorylation but met-enkephalin only showed biphasic kinetics.

**Figure 13:**
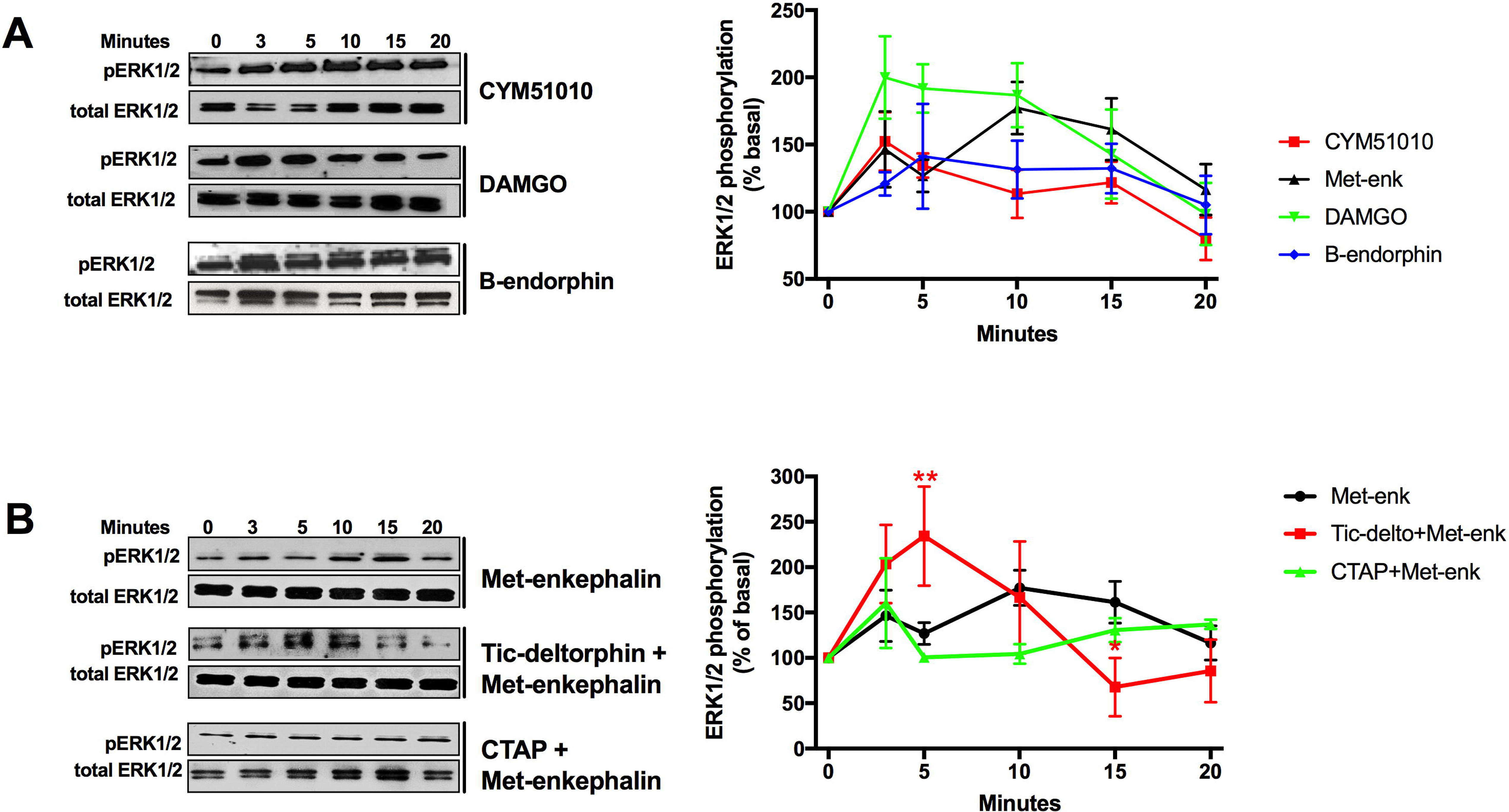
Mu-delta heteromerization alters met-enkephalin induced kinetics of ERK1/2 phosphorylation in primary hippocampal cultures. **A)** Primary hippocampal cultures were treated with DAMGO (1µM), CYM51010 (400nM), Met-enkephalin (100nM) or β-endorphin (100nM) for the indicated time. Representative Western blot with phosphorylated ERK1/2 (pERK1/2) and total ERK1/2 antibody (left panel) and quantification (right panel). At least three independent experiments were performed for each condition. **B)** Primary hippocampal cultures were treated with met-enkephalin (100nM) alone or in combination with CTAP (200nM) or Tic-deltorphin (200nM) for the indicated time. Representative Western blot with phosphorylated ERK1/2 (pERK1/2) and total ERK1/2 antibody (left panel) and quantification from at least three independent experiments were performed for each condition (right panel). Two-way ANOVA F_time_ (5, 41)=2.844, p-value=0.27. F_treatment_ (2, 41)=0.844, p= 0.438; F_interaction_ (10, 41)=2.344, p =0.271. Tukey post *hoc* test. ** p =0.0082, * p =0.029.

We next verified that the strong long-lasting ERK1/2 phosphorylation was indeed specific to mu-delta heteromer activation by pretreating the culture with the mu antagonist CTAP (200nM) or the delta antagonist naltrindole (200nM). As expected, only transient phosphorylation was observed in both cases (**Fig. 13B**). In addition, pre-treatment with naltrindole increased the level of phosphorylation (about 130%) at the short time point (**Figure 13B**).

Together, our data highlight sustained ERK1/2 phosphorylation by mu-delta heteromers in response to met-enkephalin and suggest spatio-temporal alteration in mu opioid receptor signaling.

### Mu-delta heteromerization is increased in the RVM of neuropathic animals

We have previously identified mu-delta heteromers in the RVM using mu-mCherry/delta-eGFP double fluorescent mice (Erbs et al. 2015). Since this region plays a key role in the control of the nociceptive descending pathway and is enkephalinergic (see also **Figure S4**), we examined whether the expression mu-delta heteromers is changed in neuropathic pain conditions induced by sciatic nerve injury. Indeed, the number of neurons co-expressing mu-delta almost doubled in neuropathic pain animals (91 ± 9 vs 47 ± 4 neurons/mm^2^, unpaired Student’s t test, p value=0.0097) (**Figure 14**) pointing to potential mu-delta modulation of the nociceptive response in this pathological condition.

**Figure 14:**
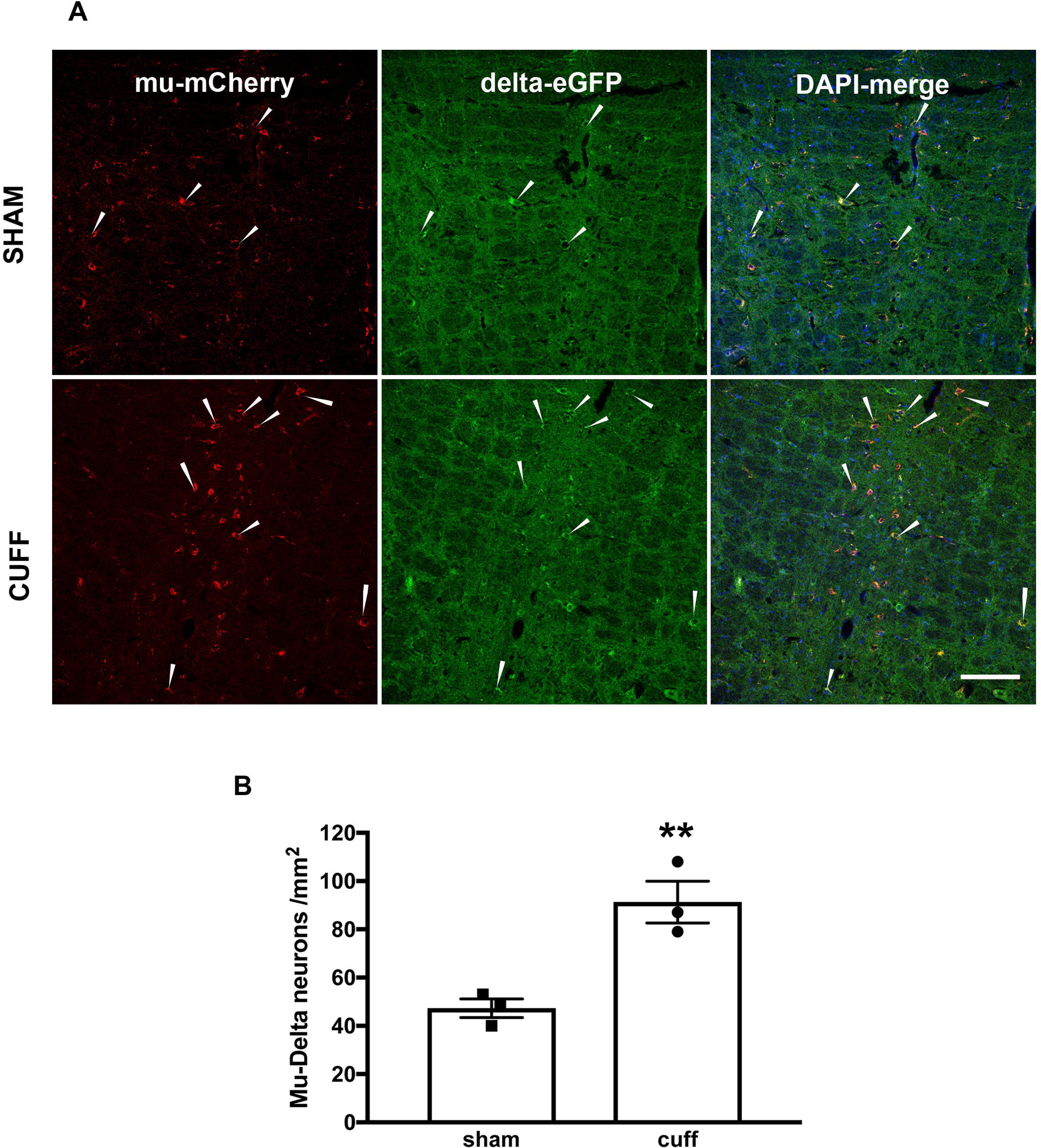
Mu-delta heteromer expression is increased in the rostral ventromedial medulla in neuropathic conditions. **A)** Representative confocal images showing mu-mCherry and delta-eGFP neuronal co-localization in the RVM in sham and neuropathic conditions. Scale bar= 50µm **B)** Quantification of the density of neurons co-expressing mu and delta opioid receptors. N=3 mice par condition. Unpaired Student’s t test, **p value=0.0097.

## Discussion

Using double fluorescent knock-in mice, we investigated the impact of heteromerization on the internalization of endogenous mu and delta opioid receptors. We first established ligand selective co-internalization upon activation by CYM51010, DAMGO and deltorphin II but not SNC80. We then evidenced mu-delta co-internalization upon activation by the endogenous opioid peptide met-enkephalin but not β-endorphin. Co-internalization was driven by the delta opioid receptor, required an active conformation of both mu and delta opioid receptors and led to sorting to the lysosomal compartment. Altered mu opioid receptor intracellular fate and sustained ERK1/2 phosphorylation were observed in response to metenkephalin. Moreover, increased expression of mu-delta heteromers was detected in the RVM under neuropathic conditions, strongly suggesting a role in the control of the descending nociceptive pathway.

### Internalization of endogenous mu-delta heteromers is ligand selective

In primary hippocampal cultures, endogenous mu-delta opioid receptors co-internalized in response to CYM51010, DAMGO or deltorphin II, but not in response to SNC80 indicating ligand specificity. In co-transfected HEK293 cells, co-internalization of mu and delta opioid receptors was reported following activation by the mu agonists DAMGO (Hasbi et al. 2007, Kabli et al. 2010, He et al. 2011) or methadone (Milan-Lobo and Whistler 2011) or following activation by the delta agonists SNC80 (Kabli et al. 2010, He et al. 2011), UFP512 (Kabli et al. 2010), deltorphin I (He et al. 2011), deltorphin II (Hasbi et al. 2007, Kabli et al. 2010, He et al. 2011) or DPDPE (Kabli et al. 2010). Differences between native and heterologously expressed receptors may reflect distinct cellular contents (Benredjem et al. 2017). Indeed, no mu-delta receptor co-internalization is detected in the spinal cord following SNC80 (10mg/kg i.p.) administration (Wang et al. 2018). Also, internalization of the delta opioid receptor is dependent on GRK2 in cortical neurons but not in transfected HEK 293 cells although the latter express GRK2 and support GRK2 mediated internalization of other GPCRs (Charfi et al. 2014). Similarly, the ability of ligands to differentially activate signaling pathways in AtT20 neuroblastoma and CHO cell lines uncovers clear influence of the cellular background on mu opioid receptor signaling (Thompson et al. 2016). Moreover, mu opioid receptor internalization in response to morphine only takes place in selected neuronal types indicating that variations among neuronal populations also exist. (Haberstock-Debic et al. 2005). Finally, high expression levels of a GPCR in a non-native environment can also artificially elicit interactions that would not occur *in vivo* and subsequently affect functional responses.

A potential influence of the fluorescent tag on the receptor subcellular distribution is also to be considered. In particular, strong surface expression of the delta-eGFP construct in the hippocampus could alter receptor trafficking and signaling. However, no overt change in the neuroanatomical distribution, pharmacological and signaling properties or behavioral response has been evidenced so far in the knock-in mice expressing the delta-eGFP and/or mu-mcherry fluorescent fusions (reviewed in (Ceredig and Massotte 2014). Importantly, delta-eGFP surface expression varies across the nervous system and is increased upon chronic morphine administration (Erbs et al. 2016) or in neuropathic pain conditions (Ceredig et al. 2018) as previously reported for wild type receptors (reviewed in (Gendron et al. 2015), (Ceredig and Massotte 2014). Moreover, the use of the delta-eGFP fusion enabled to detect *in vivo* partial receptor internalization in response to a physiological stimulation (Faget et al. 2012). The fluorescent knock-in mice therefore appear well suited reporters for native opioid receptor studies.

### Internalization of endogenous mu-delta heteromers requires both receptors in an active conformation

Phosphorylation of the delta opioid receptor C–terminus is important for agonist-induced β-arrestin 2 recruitment (Qiu et al. 2007) and mu opioid receptor phosphorylation facilitates endocytosis through enhanced receptor affinity for β-arrestin (Qiu et al. 2003). In co-transfected HEK293 cells, mu-delta receptor co-internalization by the delta agonists SNC 80 and deltorphin I or the mu agonist DAMGO is correlated to increased phosphorylation at the delta or the mu opioid receptor respectively (He et al. 2011). Moreover, the mu opioid receptor is not internalized following stimulation by the delta selective agonists SNC 80 or deltorphin I when co-expressed with a phosphorylation deficient delta opioid receptor (He et al. 2011). These results indicate that activation of the delta opioid receptor is required to promote co-internalization of the mu opioid receptor by delta agonists. Here, we show that both mu and delta opioid receptors are phosphorylated when activated by the biased mu-delta agonist CYM51010 or by the endogenous opioid peptide met-enkephalin. Both ligands promote phosphorylation at the serine S363 residue which is required for delta opioid receptor internalization by exogenous agonists in HEK293 cells (Kouhen et al. 2000, Law et al. 2000) and is also phosphorylated *in vivo* upon activation with met-enkephalin (Faget et al. 2012). Both ligands also increase phosphorylation at serine S375 and threonine T370, T376 and T379 residues necessary for mu opioid receptor internalization by exogenous ligands such as DAMGO and fentanyl or by the endogenous opioid peptide met-enkephalin (Just et al. 2013, Yousuf et al. 2015). Endogenous mu-delta receptor co-internalization is blocked by mu (CTAP, cyprodime, β-FNA) or delta (naltrindole, tic-deltorphin) selective antagonists. In cotransfected HEK 293 cells, the mu agonist CTOP blocked mu-delta receptor co-internalization by the delta agonists UFP512, SNC80 or deltorphin II whereas the delta antagonist naltrindole reduced mu-delta receptor co-internalization by the mu agonist DAMGO (Kabli et al. 2010). Moreover, internalization of the mu opioid receptor was not observed upon mu-delta receptor co-transfection in the absence of a ligand despite detection of delta constitutive internalization (Law et al. 2005). In addition, pre-treatment with pertussis toxin in primary hippocampal neurons blocked DAMGO induced mu-delta co-internalization supporting the need for a G protein activatable conformation of the mu opioid receptor. Co-internalization thus requires that both mu and delta opioid receptors are in an active conformation.

Interestingly, delta antagonists blocked mu-delta receptor co-internalization whereas mu antagonists only blocked mu opioid receptor internalization without affecting delta opioid receptor internalization and degradation. These data indicate that the delta opioid receptor is driving the co-internalization, possibly through constitutive interaction with β-arrestin (Law et al. 2000, Bradbury et al. 2009). Delta antagonists could therefore inhibit co-sequestration of the receptors by destabilizing this interaction and inducing dissociation from the heteromer as previously proposed (Rozenfeld and Devi 2007).

### The intracellular fate of mu opioid receptors is altered by mu-delta heteromerization in vivo

Mu opioid receptors rapidly recycle back to the plasma membrane *in vivo* (Trafton et al. 2000). Endogenous delta opioid receptors, in contrast are slow recycling receptors that are degraded in the lysosomal compartments (Pradhan et al. 2009). Here, we observe that endogenous mu-delta heteromers are targeted to the lysosomal compartment in primary neurons and *in vivo* triggering a change in the mu opioid receptor intracellular fate following activation by exogenous or endogenous ligands. Noteworthy, this change does not correlate with differences in the receptor phosphorylation profile suggesting that it does not determine receptor recycling or degradation. Ubiquitination controls the endocytic trafficking and sorting from the endosomal membrane to the luminal compartment in multivesicular bodies for a number of GPCRs (Skieterska et al. 2017). In co-transfected HEK293 cells, He et al (He et al. 2011) correlated deltorphin I ability to increase receptor ubiquitination with enhanced mu-delta receptor co-localization in the lysosomal compartment. However, this modification is not essential to determine the delta opioid receptor intracellular fate (Hislop et al. 2009, Henry et al. 2011) and is not involved in the targeting of mu opioid receptors to the recycling pathway though it is required for mu opioid receptor lysosomal down-regulation by ESCRT (endosomal sorting complex required for transport) (Henry et al. 2011). Mu opioid receptor endocytic trafficking between recycling and lysosomal pathways is rather regulated by the C-terminal sequence LENLEAE called mu opioid regulatory sequence (MRS) (Tanowitz and von Zastrow 2003). Accordingly, mu opioid receptor splice variants lacking this sequence fail to rapidly recycle back to the plasma membrane (Tanowitz et al. 2008). Interactions between the distal C-tails of mu and delta opioid receptors are important for mu-delta heteromerization since reduced mu-delta co-immunoprecipitation is observed in co-transfected Cos cells upon truncation of the last 15 C-terminal amino acids of the delta opioid receptor (Fan et al. 2005). In addition, reduced co-immunoprecipitation is also observed *in vivo* in the presence of an interfering peptide corresponding to the delta distal C-terminal tail (Kabli et al. 2013). Mu-delta heteromerization may therefore masks mu MRS, allowing mu opioid receptor sorting to the lysosomal compartment.

### Activation of endogenous mu-delta heteromers promotes sustained ERK1/2 phosphorylation

G protein dependent ERK 1/2 phosphorylation is mediated by βγ dimers and takes place shortly after agonist activation (Carmona-Rosas et al. 2018). Accordingly, activation of the mu opioid receptor by DAMGO promotes transient pertussis toxin sensitive ERK 1/2 phosphorylation in transfected HEK293 cells (Massotte et al. 2002). In primary hippocampal neurons, we observed sustained ERK 1/2 phosphorylation following met-enkephalin activation of mu-delta heteromers. Pre-treatment with an antagonist for either receptor abolished prolonged phosphorylation and receptor co-internalization, strongly supporting mu-delta specificity and β-arrestin 2 dependent intracellular signalling as described for DAMGO (Rozenfeld and Devi 2007). Importantly, rapid transient ERK1/2 phosphorylation by metenkephalin is increased upon pre-treatment with a delta antagonist revealing that heteromerization dampens mu opioid receptor plasma membrane associated ERK1/2 phosphorylation by promoting receptor sequestration (Weinberg et al. 2017). This in turn may trigger significant changes in the functional response such as reduced nuclear transcriptional activation (Eichel and von Zastrow 2018) or decreased activity of GIRK channels (Nagi and Pineyro 2014). In the ventral tegmental area where the two receptors functionally interact, increased G protein dependent K^+^ conductance in response to DPDPE or deltorphin II was observed in the presence of the mu antagonist CTAP. Conversely the delta antagonist TIPPψ enhanced DAMGO evoked responses (Margolis et al. 2017) suggesting that mu-delta functional interactions blunt this signalling pathway *in vivo*.

Interestingly, heteromerization of the mu opioid and somatostatin sst2 receptors also induces sustained β-arrestin 2 dependent ERK1/2 phosphorylation in pancreatic ductal adenocarcinoma (Jorand et al. 2016). However, heteromerization of the mu and galanin1 receptors generates bidirectional negative cross-talk on ERK phosphorylation (Moreno et al. 2017), indicating that modulation of the ERK pathway varies along with the receptor pair. The spatiotemporal control of mu opioid receptor signalling via heteromerization with the delta opioid receptor therefore reveals a specific mechanism by which the endogenous opioid system can modulate neuronal activity.

### In vivo functional consequences of mu-delta heteromerization in the RVM

Association with the delta opioid receptor alters the spatio-temporal dynamics of the mu opioid receptor when activated by met-enkephalin and questions the physiological role of mu-delta heteromers *in vivo*. Enhanced mu-dependent surface expression of the delta opioid receptor in the spinal cord and the dorsal root ganglia is triggered by neuropathic conditions ((Morinville et al. 2003, Gendron et al. 2006), reviewed in (Gendron et al. 2015)). Here, we observed increased mu-delta heteromer expression in the RVM following sciatic nerve cuffing that reflects supraspinal adaptation to the pathological condition. Co-expression of functional mu and delta opioid receptors in this structure has been described in GABAergic interneurons that modulate the descending nociceptive pathway (Pedersen et al. 2011). Mu-delta targeting to the lysosomal compartment and blunted plasma membrane associated ERK1/2 phosphorylation point to decreased mu opioid receptor signalling that would reduce the GABAergic inhibitory tonus exerted on the descending nociceptive pathway and suggest an antinociceptive role for mu-delta heteromers.

In a rat model of CFA, functional synergy was observed between DAMGO and the delta agonist deltorphin II though no change in receptor expression, affinity or potency could be detected in the RVM (Sykes et al. 2007). This synergic effect could be explained by the positive cooperativity observed in ligand binding to mu-delta heteromers (Gomes et al. 2011) since enkephalins have slightly better affinity for delta opioid receptors (Lord et al. 1977). Several lines of evidence indicate a critical role for enkephalin in the RVM besides the well-established role of peripheral enkephalins in the context of chronic pain (Lesniak and Lipkowski 2011). Chronic inflammatory conditions enhance enkephalin expression in the RVM (Hurley and Hammond 2001) and microinjection of enkephalin in this structure is antinociceptive in a model of inflammatory pain (Little et al. 2012). Also, exercise decreases neuropathic pain following sciatic nerve ligation in rat by increasing enkephalin levels in the RVM (Stagg et al. 2011) and i.c.v. injection of the delta antagonist BNTX suppresses enkephalin-mediated antinociception (Takemori and Portoghese 1993). Importantly, i.c.v. injection of the delta antagonists BNTX or naltriben do not impact β-endorphin antinociception suggesting no involvement of the delta opioid receptor in β-endorphin mediated effects (Takemori and Portoghese 1993). These findings are in full agreement with our observation that enkephalin but not β-endorphin alters mu-delta heteromer function.

In conclusion, our data demonstrate that mu-delta heteromerization alters the functional dynamics of the mu opioid receptor physiological response which provides a way to fine-tune mu opioid receptor signalling in pathological conditions. It also represents an interesting emerging concept for the development of novel therapeutic drugs and strategies.

## Acknowledgments

The authors would like to thank the Chronobiotron animal facility (UMS 3415 CNRS), and the in vitro imaging platform of the Institut des Neurosciences Cellulaires et Intégratives (UPS 3156 CNRS) for their assistance. The authors would also like to acknowledge Drs C.M. Cahill and I. Gomes for critical reading of the manuscript, and Elisabeth Waltisperger for excellent technical contribution.

The work was performed thanks to the financial support of the Fondation pour la Recherche Médicale (DPA20140129364), the CNRS and the University of Strasbourg and, the Interdisziplinäres Zentrum für klinische Forschung (Juniorprojekt J61). L. Derouiche was the recipient of an IDEX post-doctoral fellowship of the University of Strasbourg. M. Ugur was a fellow of the Erasmus Mundus Joint PhD program Neurotime.

## Author contributions

LD, DM, SS designed experiments

LD, MU, FP, AM, SD, DM, SO conducted experiments and analysed data

LD, DM, SO wrote the manuscript

## Declaration of interest

The authors declare no competing interests.

